# Fine-mapping, trans-ancestral and genomic analyses identify causal variants, cells, genes and drug targets for type 1 diabetes

**DOI:** 10.1101/2020.06.19.158071

**Authors:** C.C. Robertson, J.R.J. Inshaw, S. Onengut-Gumuscu, W.M. Chen, D. Flores Santa Cruz, H. Yang, A.J. Cutler, D.J.M. Crouch, E. Farber, S.L. Bridges, J.C. Edberg, R.P. Kimberly, J.H. Buckner, P. Deloukas, J. Divers, D. Dabelea, J.M. Lawrence, S. Marcovina, A.S. Shah, C.J. Greenbaum, M.A. Atkinson, P.K. Gregersen, J.R. Oksenberg, F. Pociot, M.J. Rewers, A.K. Steck, D.B. Dunger, Type 1 Diabetes Genetics Consortium, L.S. Wicker, P. Concannon, J.A. Todd, S.S. Rich

**Author notes:** These authors contributed equally to this work (C.C. Robertson, J.R.J. Inshaw). These authors jointly supervised this work (J.A. Todd, S.S. Rich). Address Correspondence to: John A Todd, FRS, FMedSci, PhD; Professor of Precision Medicine; Director of the Wellcome Centre for Human Genetics and the JDRF/Wellcome Diabetes and Inflammation Laboratory; Wellcome Centre for Human Genetics; University of Oxford; Roosevelt Drive, Oxford, OX3 7BN, United Kingdom.

## Abstract

We report the largest and most ancestrally diverse genetic study of type 1 diabetes (T1D) to date (61,427 participants), yielding 152 regions associated to false discovery rate < 0.01, including 36 regions associated to genome-wide significance for the first time. Credible sets of disease-associated variants are specifically enriched in immune cell accessible chromatin, particularly in CD4^+^ effector T cells. Colocalization with chromatin accessibility quantitative trait loci (QTL) in CD4^+^ T cells identified five regions where differences in T1D risk and chromatin accessibility are potentially driven by the same causal variant. Allele-specific chromatin accessibility further refined the set of putative causal variants with functional relevance in CD4^+^ T cells and integration of whole blood expression QTLs identified candidate T1D genes, providing high-yield targets for mechanistic follow-up. We highlight rs72938038 in *BACH2* as a candidate causal T1D variant, where the T1D risk allele leads to decreased enhancer accessibility and *BACH2* expression in T cells. Finally, we prioritise potential drug targets by integrating genetic evidence, functional genomic maps, and immune protein-protein interactions, identifying 12 genes implicated in T1D that have been targeted in clinical trials for autoimmune diseases. These findings provide an expanded genomic landscape for T1D, including proposed genetic regulatory mechanisms of T1D-associated variants and genetic support for therapeutic targets for immune intervention.

## Main

Type 1 diabetes (T1D) is characterized by an autoimmune attack that destroys the insulin-producing pancreatic β cells and is driven by diverse genetic^1–6^ and environmental^7^ factors. Approximately 50 chromosome regions are known to contain variants that alter T1D risk^1–6^, substantially fewer than other common diseases. Less is known about the contribution of T1D risk alleles in non-European populations, despite recent increases in T1D diagnoses in multiple non-European ancestry groups^8^. To address these gaps in T1D genetics, we doubled the sample size from the previous largest T1D association study, genotyped ancestrally diverse T1D cases, controls, and affected families with the Illumina ImmunoChip^3^ array, and imputed additional variants using a large haplotype reference panel^9^.

Genetic screening and autoantibody surveillance can detect islet autoimmunity before overt progression to diabetes^10,11,12^, providing an opportunity for prevention in T1D. However, there are currently no interventions that prevent T1D. With the understanding that T1D is an immune-mediated disease, multiple immune therapies have been explored in clinical trials, with some altering disease course^13^. In a recent clinical trial, a 14-day course of teplizumab, an anti-CD3 monoclonal antibody, was shown to delay T1D diagnosis in high-risk individuals by a median of two years^14^ This success provides evidence that appropriately-timed immune-modulating therapy can alter the autoimmune process preceding clinical disease. More precise targets and better characterization of their role in disease initiation and progression will provide opportunities for safe and effective intervention and, ultimately, prevention of T1D. Drug targets based on genetic evidence are more likely to succeed in clinical trials^15,16^. Thus, carefully defining the genetic factors contributing to T1D risk and how they disrupt immune pathways in affected individuals can help guide development and repurposing of immune-modulating therapies for T1D prevention.

Fine-mapping genetic regions is an important step towards understanding the mechanism of a genetic association, since it clarifies the number of independent disease associations in each region and identifies the most likely causal variants, or ‘credible variants’, for each association. Differences in linkage disequilibrium (LD) structure between ancestry groups can be advantageous in prioritising causal variants, since only a portion of a large haplotype in one ancestry group might be associated with disease in a different ancestry group. We showed previously that T1D credible variants are most strongly enriched in lymphocyte and thymic enhancers^3^. Integration of fine-mapped T1D risk variants and cell type-specific expression and chromatin accessibility quantitative trait loci (eQTL and caQTL) data provide a means to assess whether the variant altering disease risk is also altering gene expression or chromatin activity, helping to prioritise variants and genes for interrogation of molecular mechanisms underlying disease association. Further integration of genes implicated by fine-mapping and molecular maps with immune protein networks can define a set of prioritised targets for pharmacologic intervention^17^.

Here, we define the genetic landscape of T1D and the functional impact of the variants using eQTL and caQTL data, highlighting a compelling hypothesis of genetic regulatory mechanism in the T1D locus containing the transcription factor BACH2. In addition, we highlight drugs that target T1D gene candidates or networks that they act within, which may impact and inform strategies for treatment or prevention in T1D.

## Results

### Thirty-six new regions at p < 5×10^-8^

After quality filtering, 61,427 participants (**Supplementary Table 1**) and 140,333 genotyped ImmunoChip variants (Online Methods) were included in analysis, providing dense coverage in 188 autosomal regions (“ImmunoChip regions”)^18^ and sparse genotyping in other regions of the genome (**Supplementary Tables 2 and 3**). Each participant was assigned to one of five ancestry groups using k-means clustering of ImmunoChip genotype principal components (Online Methods, **Supplementary Figure 1**) - European (EUR, N = 47,319), African-admixed (AFR, N = 4,290), Finnish (FIN, N = 6,991), East Asian (EAS, N = 588) and Admixed (AMR, N = 2,239). In total, 16,159 T1D cases, 25,386 controls and 6,143 trio families (i.e., an affected child and both parents) were genotyped and included in association analyses (**Supplementary Tables 4 and 5**). Imputation with the National Heart, Lung, and Blood Institute (NHLBI) Trans-Omics for Precision Medicine (TOPMed)^9^ multi-ethnic reference panel was used to improve discovery power and fine-mapping resolution (Online Methods). After imputation, the number of variants in ImmunoChip regions with imputation R-squared > 0.8 and MAF > 0.005 in each ancestry group was 322,084 (AFR), 166,274 (EUR), 163,612 (FIN), 137,730 (EAS) and 188,550 (AMR). We compared imputed genotypes to whole genome sequencing data from a subset of individuals, with high concordance observed (Online Methods, **Supplementary Note 1**).

Initially, we analyzed unrelated cases and controls (N = 41,545), assuming an additive inheritance model. With minimal evidence of artificial inflation of association statistics due to population structure (**Supplementary Note 2**), we identified 64 T1D-associated regions outside the major histocompatibility complex (MHC) HLA region, including 24 regions associated with T1D at genome-wide significance for the first time (p < 5×10^-8^). One of these associations was proximal to the gene encoding the interleukin-6 receptor (IL-6R). However, despite both variants being directly genotyped, the lead variant in this region was rs2229238 (C>T, p = 3.02×10^-9^), not the nonsynonymous variant rs2228145 (A>C; *IL6R* Asp358Ala; p = 2.20×10^-4^), which is associated with RA^19^ and previously suggested to be causal for T1D^20^. Following conditional analysis, 78 independent associations were identified (p < 5×10^-8^; **Supplementary Table 6**). No genome-wide associations were identified on the X chromosome, where the most associated variant was rs4326559 (A>C, C allele OR = 1.09, p = 4.5×10^-7^).

We extended the discovery analysis to incorporate T1D trio families (N = 6,143, some trio families were multiplex but analyzed as trios, Online Methods). Meta-analysis of case-control and trio results identified 78 chromosome regions associated with T1D (p < 5×10^-8^), including 42/43 chromosome regions previously identified in an ImmunoChip-based study^3^ (rs4849135 (G>T) was p = 2.93×10^-7^) and 36 novel regions associated with T1D at genome-wide significance for the first time (**Table 1**).

**Table 1:** Regions of association with T1D, identified to genome-wide significance (p < 5×10^-8^) for the first time. Of these 36 regions, 13 had a lead variant that was in linkage disequilibrium (r^2^ > 0.95 in 1000 Genomes Project European population) with variants that are associated with at least one other related trait.

| Chr | Position (bp) † | Lead Variant rsID | A1 | A2 | Putative candidate gene* | AF<sub>EUR</sub> (A2) | OR**<sub>META</sub> | P<sub>META</sub> | Other related trait associations*** |
| --- | --- | --- | --- | --- | --- | --- | --- | --- | --- |
| 1 | 63643100 | rs2269241 | T | C | *PGM1* | 0.196 | 1.111 | $4.67 \times 10^{-12}$ | |
| 1 | 92358141 | rs34090353 | G | C | *RPAP2* | 0.361 | 1.078 | $1.10 \times 10^{-8}$ | |
| 1 | 119895261 | rs2641348 | A | G | *NOTCH2* | 0.107 | 1.113 | $1.61 \times 10^{-8}$ | Crohn's, T2D |
| 1 | 154465420 | rs2229238 | T | C | *IL6R* | 0.813 | 0.896 | $1.38 \times 10^{-12}$ | |
| 1 | 172746562 | rs78037977 | A | G | *FASLG* | 0.124 | 0.884 | $2.41 \times 10^{-9}$ | Asthma, vitiligo, allergic sensitization |
| 1 | 192570207 | rs2816313 | G | A | *RGS1* | 0.719 | 1.090 | $4.57 \times 10^{-9}$ | |
| 1 | 212796238 | rs11120029 | G | T | *TATDN3* | 0.147 | 1.102 | $1.82 \times 10^{-8}$ | |
| 2 | 12512805 | rs10169963 | C | T | *AC096559.1* | 0.580 | 1.074 | $2.78 \times 10^{-8}$ | |
| 2 | 100147438 | rs12712067 | G | T | *AFF3* | 0.358 | 0.925 | 4.12×10<sup>-9</sup> |  |
| 2 | 191105394 | rs7582694 | C | G | *STAT4* | 0.773 | 0.916 | 2.83×10<sup>-9</sup> | SLE, hypothyroidism, coeliac, RA |
| 2 | 241468331 | rs10933559 | A | G | *FARP2* | 0.208 | 1.109 | 2.39×10<sup>-11</sup> |  |
| 4 | 973543 | rs113881148 | C | A | *TMEM175* | 0.626 | 1.082 | 5.72×10<sup>-9</sup> | Body fat percentage |
| 4 | 38602849 | rs337637 | G | A | *KLF3* | 0.364 | 0.919 | 2.57×10<sup>-10</sup> | White blood cell count |
| 5 | 40521603 | rs1876142 | G | T | *PTGER4* | 0.658 | 0.905 | 2.18×10<sup>-14</sup> |  |
| 5 | 56146422 | rs10213692 | T | C | *ANKRD55/IL6ST* | 0.241 | 0.912 | 2.85×10<sup>-9</sup> | RA, Crohn's, MS |
| 6 | 424915 | rs9405661 | C | A | *IRF4* | 0.514 | 1.080 | 2.26×10<sup>-9</sup> |  |
| 6 | 137682468 | rs12665429 | T | C | *TNFAIP3* | 0.370 | 0.907 | 1.36×10<sup>-13</sup> |  |
| 6 | 159049210 | rs212408 | G | T | *TAGAP* | 0.638 | 1.112 | 1.42×10<sup>-15</sup> | MS, Crohn's, eczema |
| 7 | 20557306 | rs17143056 | A | G | *ABCB5* | 0.183 | 0.909 | 2.44×10<sup>-8</sup> |  |
| 7 | 28102567 | rs10245867 | G | T | *JAZF1* | 0.331 | 0.928 | 3.15×10<sup>-8</sup> | Eczema, hay fever, MS, SLE, monocyte percentage |
| 8 | 11877675 | rs2250903 | G | T | *CTSB* | 0.283 | 0.905 | 1.35×10<sup>-10</sup> |  |
| 9 | 99823263 | rs1405209 | T | C | *NR4A3* | 0.375 | 1.075 | 3.45×10<sup>-8</sup> |  |
| 10 | 33137219 | rs722988 | T | C | *NRP1* | 0.367 | 1.108 | 3.21×10<sup>-15</sup> |  |
| 11 | 35267496 | rs11033048 | C | T | *SLC1A2* | 0.366 | 1.091 | 1.53×10<sup>-10</sup> | Vitiligo |
| 11 | 60961822 | rs79538630 | G | T | *CD5/CD6* | 0.035 | 1.213 | 1.14×10<sup>-9</sup> |  |
| 11 | 61828092 | rs968567 | C | T | *FADS2* | 0.177 | 0.903 | 8.42×10<sup>-9</sup> | RA, neutrophil percentage |
| 11 | 64367826 | rs645078 | A | C | *CCDC88B* | 0.385 | 0.925 | 3.34×10<sup>-9</sup> |  |
| 11 | 128734337 | rs605093 | G | T | *FLII* | 0.470 | 1.077 | 4.25×10<sup>-9</sup> |  |
| 12 | 8942630 | rs1805731 | T | C | *M6PR* | 0.389 | 1.073 | 4.16×10<sup>-8</sup> | Eosinophil count |
| 12 | 53077434 | rs7313065 | C | A | *ITGB7* | 0.162 | 1.101 | 3.28×10<sup>-9</sup> |  |
| 13 | 42343795 | rs74537115 | C | T | *AKAP11* | 0.141 | 1.109 | 5.41×10<sup>-9</sup> |  |
| 14 | 68286876 | rs911263 | C | T | *RAD51B* | 0.710 | 1.083 | 1.69×10<sup>-8</sup> | PBC, SLE, RA |
| 16 | 20331769 | rs4238595 | T | C | *UMOD* | 0.687 | 0.912 | 2.43×10<sup>-11</sup> |  |
| 17 | 45996523 | rs1052553 | A | G | *MAPT* | 0.232 | 0.879 | 1.65×10<sup>-15</sup> | Parkinson's disease |
| 17 | 47956725 | rs2597169 | A | G | *PRR15L* | 0.348 | 1.081 | 3.35×10<sup>-9</sup> |  |
| 21 | 44204668 | rs56178904 | C | T | *ICOSLG* | 0.187 | 0.898 | 6.48×10<sup>-11</sup> |  |

†genome build 38
\*Closest gene or gene with mechanistic support from the literature.
\*\*Additive odds ratio for the addition of an A2 allele.
\*\*\*Variant with 1000 Genomes Project European population LD $r^2 > 0.95$ with lead variant associated with any trait
potentially relevant trait (https://genetics.opentargets.org). RA= rheumatoid arthritis, T2D= type 2 diabetes, SLE= systemic lupus erythematosus, MS= multiple sclerosis, IBD= irritable bowel disease, PBC= primary biliary cholangitis.
AF = Allele Frequency, OR= Odds Ratio

Applying Benjamini-Yekutieli FDR < 0.01^21^ to assess statistical significance, 143 regions were associated with T1D (**Supplementary Table 8;** A full set of summary statistics can be found in **Supplementary Table 9**). The absolute effect sizes for FDR < 0.01 associated variants not reaching the threshold for genome-wide significance (p < 5×10^-8^) were smaller than those satisfying genome-wide significance and with similar MAFs (median (IQR)) OR = 1.07 (1.06, 1.09) versus 1.11 (1.09, 1.13); median (IQR) MAF = 0.301 (0.152, 0.397) versus 0.306 (0.184, 0.374)). These results indicate that there may be many more regions associated with T1D with increasingly smaller effect sizes (**Supplementary Figure 4**), requiring genome-wide coverage and larger sample sizes for detection.

One exception was chromosome 1p22.1 near the Metal Response Element Binding Transcription Factor 2 (MTF2) gene, where the minor allele (A) at the lead variant, rs190514104 (G>A), is rare in most ancestry groups (< 0.1%), but more common in the AFR ancestry group (> 1%) and had a large effect on T1D risk (OR (95% CI) = 2.9 (1.9-4.5); p = 6.6×10^-7^). This example illustrates how focusing on previously understudied ancestry groups in future T1D studies could lead to more biological insight, even with limited sample sizes.

Use of recessive and dominant models of inheritance identified 35 regions (25 dominant, 10 recessive) with a better fit than the additive model (lower Akaike Information Criterion (AIC) in Europeans) at FDR < 0.01, including nine new regions that did not reach FDR < 0.01 under the additive model (**Supplementary Table 7**). Thus, a total of 152 regions were associated with T1D at FDR < 0.01, 143 under an additive model and nine under recessive or dominant models.

### Fine-mapping reveals over one third of T1D loci contain more than one independent association

To define the local architecture of T1D regions, we applied a Bayesian stochastic search method (GUESSFM^22^) in ImmunoChip regions only (Online Methods, **Supplementary Table 2**), to identify credible variants in the European case-control data (Online Methods). Of 52 ImmunoChip regions associated with T1D, 21 (40%) were predicted to contain more than one causal variant (**Figure 1a**), compared to nine regions with evidence of multiple association signals using stepwise conditional regression. Moreover, in four regions identified using a conventional stepwise logistic regression approach, the lead variant in the discovery analysis was not prioritised by fine-mapping (posterior probability < 0.5): 2q33.2 (*CTLA4*), 4q27 (*IL2*), 14q32.2 (*MEG3*) and 21q22.3 (*UBASH3A*). In these regions, the lead variant likely tags two or more disease-associated haplotypes that can be identified using GUESSFM but not stepwise logistic regression, a phenomenon observed previously^22,23^. To illustrate this, stepwise logistic regression analysis of the *UBASH3A* locus supports a single causal variant, rs11203203 (G>A) (**Supplementary Table 6**), while GUESSFM fine-mapping supports a three-variant model (rs9984852 (T>C), rs13048049 (G>A) and rs7276555 (T>C)), each having a much weaker association with T1D when examined in univariable analyses (**Figure 1b**). However, the GUESSFM three-variant model has a better fit than the single variant model, as indicated by a lower AIC (45073 vs. 45138, **Figure 1c**). Haplotype analysis (Online Methods) demonstrated that when the risk allele for the lead variant from discovery analysis, rs11203203 (A), is present without any of the three GUESSFM-prioritised variants, there is no effect on disease risk; thus, rs11203203 is unlikely to be casual for T1D (**Figure 1d**). Given the complexity of association at many loci, statistical methods designed to use univariable summary statistics alone may not be the most effective way to explore the genetic architecture of T1D. The comprehensive list of credible variant sets is provided in **Supplementary Table 11**, and all haplotype analyses can be viewed at https://github.com/ccrobertson/t1d-immunochip-2020).

**Figure 1:**
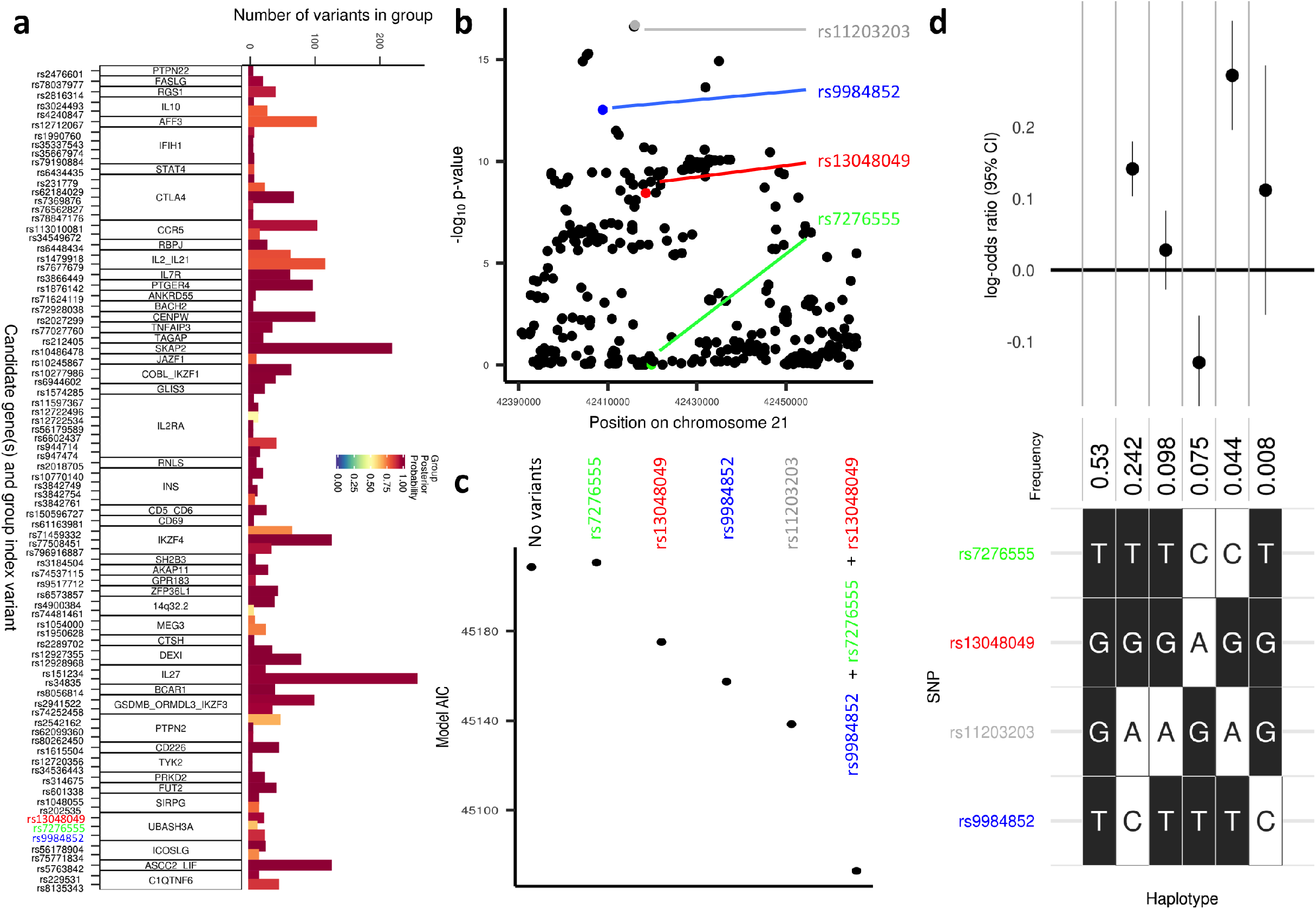
Fine-mapping T1D regions using a Bayesian stochastic search algorithm. **a)** Number of variants in GUESSFM-prioritised groups with group posterior probability > 0.5. Candidate gene names and lead variants for each group are shown on the y-axis. **b)** Manhattan plot of the *UBASH3A* region from the EUR case-control analysis, highlighting the lead variant from the univariable analysis, rs11203203 (grey) and the three variants prioritised using GUESSFM, rs9984852 (blue), rs13048049 (red) and rs7276555 (green). **c)** Comparison of model AIC in the *UBASH3A* region from a model including EUR cases and controls only, comparing combinations of alleles prioritised either in a univariable (grey) or GUESSFM analyses (red, green and blue), showing the three-variant model prioritised by GUESSFM has a lower AIC (better fit) than the single variant model from the univariable analysis. **d)** Haplotype analysis including the lead univariable variant and the three GUESSFM-prioritised variants from the *UBASH3A* region; the effect estimate from the haplotype which contains the minor allele at rs11203203 (grey) only relative to the haplotype with major alleles at each variant is close to 0 (comparison of columns 1 and 3), whereas the non-zero effects of each of rs729984852 (blue, columns 2 and 6), rs13048049 (red, column 4) and rs7276555 (green, comparing columns 3 and 5) can be observed.

In the 30 regions where our analysis suggested a single causal variant, multi-ethnic fine-mapping was performed using PAINTOR^24^. Eight regions showed an associated variant (p < 5×10^-4^) in more than one ancestry group: five with associations in EUR and FIN, and three with associations in EUR and AFR. In three chromosome regions, the number of variants prioritised was markedly reduced by including multiple ancestry groups: 4p15.2 (*RBPJ*), 6q22.32 (*CENPW*) and 18q22.2 (*CD226*) (**Figure 2a, Supplementary Figures 6 and 7, Supplementary Table 12**). At chromosome 4p15.2 (*RBPJ*), the credible set based on EUR subjects contained 24 variants, while using PAINTOR with EUR and AFR summary statistics, just five variants were prioritised with a posterior probability > 0.1 (**Figure 2a**). Among these prioritised variants located in the non-coding transcript *LINC02357*, two variants rs34185821 (A>G) and rs35944082 (A>G), have the potential to disrupt multiple transcription factor binding motifs^25^; rs35944082 also overlaps open chromatin in multiple adaptive immune cell types (**Figure 2b**) and resides in a FANTOM enhancer site (http://slidebase.binf.ku.dk/human_enhancers). Further, rs34185821 is one of three prioritised variants flanking an activation-dependent ATAC-seq peak in lymphocytes and a stable response element in human islets^26^, with potential to perturb an extended TATA box motif^27^.

**Figure 2:**
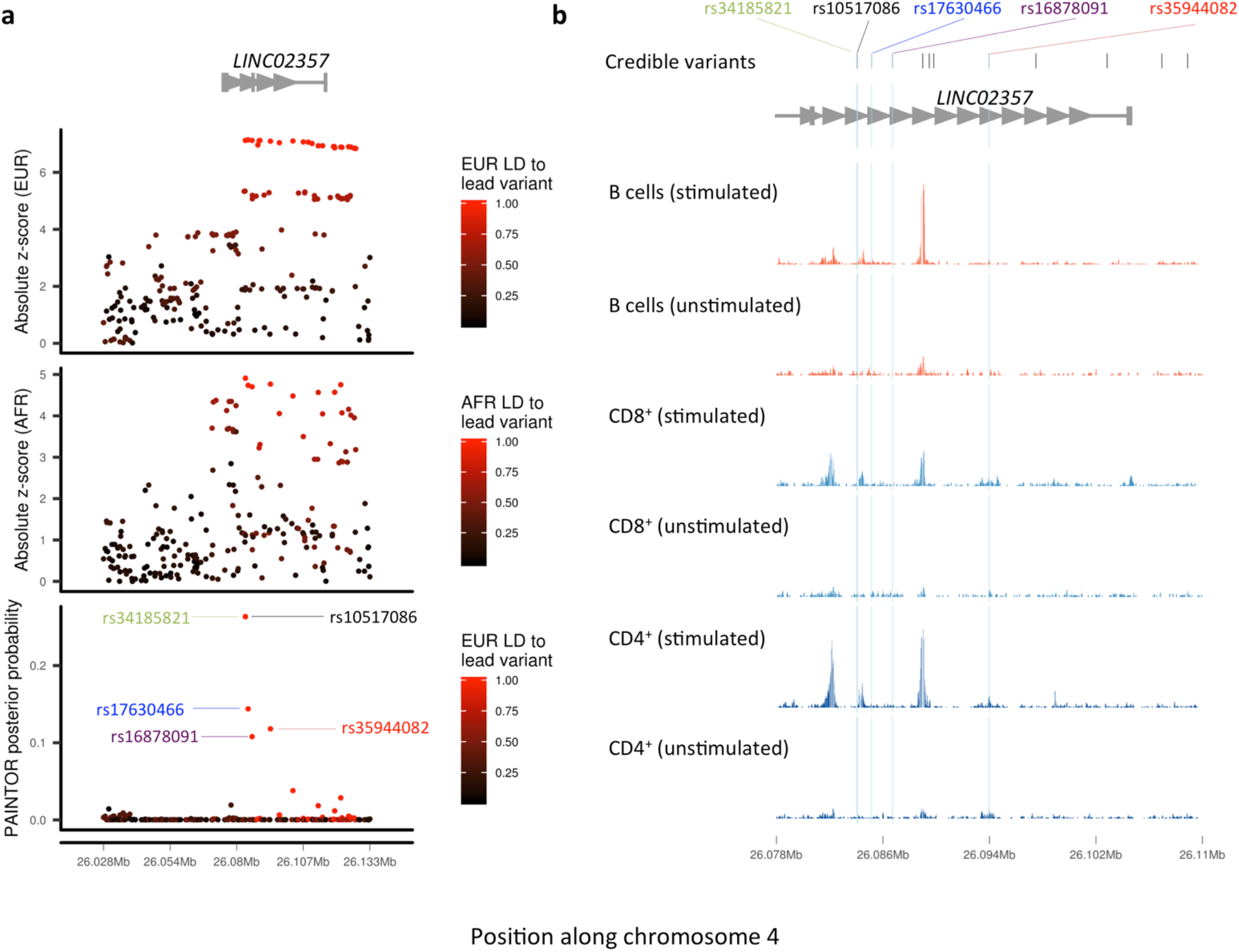
Fine-mapping of the chromosome 4p15.2 region. **a)** European (EUR, top panel) and African (AFR, middle panel) ancestry group association z-score statistics; posterior probabilities (bottom panel) from multi-ethnic fine-mapping of EUR and AFR using PAINTOR; z-scores are colored by linkage disequilibrium (LD) to the lead PAINTOR-prioritised variant. **b)** overlay of T1D-credible variants with open chromatin ATAC-seq peaks in immune cells, with variants prioritised by PAINTOR (posterior probability > 0.1) highlighted in blue. Normalized ATAC-seq read count shown for effector CD4^+^ T cells, B cells, and CD8^+^ T cells, under stimulated and nonstimulated conditions.

### T1D-associated protein-altering variants

Only 34/2732 (1.2%) credible variants (group posterior probability > 0.5) were protein-altering (nonsynonymous, frameshift, stop-gain, or splice-altering) with 12 having support for a role in T1D (Online Methods, **Supplementary Table 13**). Of note, we identified several previously unreported protein-altering variants as highly prioritised in T1D credible sets (posterior probability > 0.1): a protective missense variant in *UBASH3A*, rs13048049 (G>A; *UBASH3A* Arg324Gln; OR = 0.84; AF_EUR_ = 0.051); two low-frequency splice donor variants in *IFIH1*, rs35732034 (C>T; OR = 0.63; AF_EUR_ = 0.0089) and rs35337543 (C>G; OR = 0.61; AF_EUR_ = 0.0099); and a missense variant in *CTLA4*, rs231775 (A>G; *CTLA4* Thr17Ala; OR = 1.20; AF_EUR_ = 0.36).

### T1D credible variants are over-represented in accessible chromatin in lymphocytes, NK and dendritic cells

ATAC-seq offers a high-resolution map of accessible chromatin with regulatory function^28^. As a first step in investigating the function of T1D-associated credible variants in non-coding regions, we analyzed their overlap with accessible chromatin. Using both publicly available and newly generated chromatin accessibility (ATAC-seq) data from healthy donors^29–31^, we assessed enrichment (Online Methods) of 2,431 T1D credible variants (group posterior probability > 0.8) in accessible chromatin across diverse immune and non-immune cell types (including 25 primary immune cell types, pancreatic islets, and, as control cell types unlikely to be central to T1D etiology, fetal and adult cardiac fibroblasts). T1D credible variants were enriched in open chromatin in the majority of the primary immune cell types tested (p < 9.1 ×10^-4^). There was no enrichment in pancreatic islets (p = 0.14), the primary target of autoimmunity in T1D, _even after exposure to proinflammatory cytokines_ (p = 0.05), or in cardiac fibroblasts (p > 0.60) (**Supplementary Figure 8**).

We next examined enrichment for T1D credible variants in stimulation-responsive accessible chromatin in immune cells (Online Methods). Of 138,596 regions in the consensus list of ATAC-seq peaks, Th17 cells had the highest proportion of differentially-open peaks (FDR < 0.01) after stimulation (15.3%), while effector-memory CD8^+^ T cells had the highest proportion of differentially-open peaks in unstimulated cells (9.8%) (**Supplementary Table 14**). T1D credible variants were enriched in differentially-open chromatin after stimulation in numerous cell types, with the largest enrichment in effector CD4^+^ T cells stimulated for 24 hours with anti-CD3/CD28 and human IL-2 (**Supplementary Figure 9)**. These results indicate that T1D credible variants may contribute to islet autoimmunity, in part, by altering responses to T cell receptor signaling.

### Colocalization of T1D association with QTLs in immune cells

Chromatin accessibility profiles were generated across 115 participants (N_EUR_ = 48, N_AFR_ = 67) in primary CD4^+^ T cells, a cell type in which accessible chromatin is strongly enriched for T1D credible variants (**Supplementary Figure 8**). Additive effects of genotype on local chromatin accessibility (*cis* window < 1 Mb) were examined, identifying eleven “peaks” of chromatin accessibility significantly associated with T1D credible variants (p < 5×10^-5^). Colocalization was examined between T1D associations and caQTLs (*coloc* R package^32^, Online Methods). Five regions supported a common causal variant underlying association with T1D and chromatin accessibility (PP.H4.abf > 0.8, **Table 2**). In all five regions, at least one T1D credible variant overlapped the caQTL-associated peak. At six of these “within-peak” credible variants that were directly genotyped on the ImmunoChip, we examined allele-specific accessibility among heterozygous participants (Online Methods). The proportion of ATAC-seq reads from heterozygotes containing the alternative allele was always consistent with the direction of the caQTL effect (**Supplementary Table 15**). Within-peak credible variants with consistent caQTL effects and allele-specific accessibility provide high priority candidate variants for functional follow-up, as the observed allele-specific accessibility is unlikely to be caused by LD with variants outside the peak of interest. When integrated with whole blood *czs*-eQTLs^32,33^, colocalization identified T1D candidate genes in four of five T1D-caQTL regions (PP.H4.abf > 0.8, **Table 2**). Regions with support for a common variant underlying association with T1D, chromatin accessibility in CD4^+^ T cells, and gene expression in whole blood provide candidate causal T1D variants and genes for mechanistic investigation.

**Table 2:** T1D-associations colocalizing with caQTLs in CD4^+^ T cells. Five regions show colocalization between T1D and a caQTL with a colocalization posterior probability > 0.8. In all of these regions, at least one T1D credible variant overlaps the caQTL peak itself. In four regions, the T1D association also colocalizes with an eQTL for expression of one or more genes in whole blood.

| T1D lead variant* | Beta<sub>T1D</sub><br>** | Peak | T1D-credible<br>variants in<br>peak | caQTL lead variant* | Beta<sub>caQTL</sub><br>** | P<sub>caQTL</sub> | PP | Whole blood *cis*-eQTLs *** |
| --- | --- | --- | --- | --- | --- | --- | --- | --- |
| rs71624119<br>(chr5:56144903:G:A) | -0.099 | chr5:56147972-56149111 | rs7731626 | rs7731626<br>(chr5:56148856:G:A) | -0.5 | 2.4x10<sup>-09</sup> | 0.97 | *ANKRD55* (z=-58; PP =0.98)<br>*IL6ST* (z=-10; PP =0.98) |
| rs72928038<br>(chr6:90267049:G:A) | 0.172 | chr6:90266766-90267747 | rs72928038 | rs72928038<br>(chr6:90267049:G:A) | -1.0 | 3.9x10<sup>-16</sup> | 1.00 | *BACH2* (z=-21; PP =1) |
| rs2027299<br>(chr6:126364681:G:C) | 0.147 | chr6:126339725-126340580 | rs9388486 | rs1361262<br>(chr6:126380821:T:C) | -0.4 | 2.0x10<sup>-16</sup> | 0.87 | *CENPW* (z=-9.8; PP =0.82) |
| rs61555617<sup>†</sup><br>(chr12:56047884:TA:T) | 0.257 | chr12:56041256-56042638 | rs705704<br>rs705705 | rs705704<br>(chr12:56041628:G:A) | -0.2 | 1.1x10<sup>-15</sup> | 0.97 | *GDF11* (z=-7.5<sup>††</sup>; PP =0.97) |
| rs4900384<br>(chr14:98032614:A:G) | 0.118 | chr14:98018322-98019163 | rs11628807<br>rs4383076<br>rs11628876<br>rs11160429 | rs11628807<br>(chr14:98018774:T:G) | 0.7 | 1.8x10<sup>-21</sup> | 0.95 | - |

\* T1D lead variant is the most associated variant in the credible set, as defined by fine-mapping (Supplementary Table 11); caQTL lead variant is the most associated variant with chromatin accessibility at the peak of interest; Variants are provided as rsid (chromosome:hg38\_position:reference:alternative).
\*\* Beta<sub>T1D</sub> refers to the effect size for the alternative allele of the T1D lead variant; Beta<sub>caQTL</sub> refers to the effect size for the alternative allele of the caQTL lead variant.
\*\*\* Whole blood *cis*-eQTL statistics from eQTLGen for the T1D lead variant and colocalization with the T1D association.
<sup>†</sup> rs61555617 is referred to as rs796916887 in supplementary tables
<sup>††</sup> *cis*-eQTL statistics for rs61555617 are missing in eQTLGen; the reported *GDF11* *cis*-eQTL z-score is for the highly correlated variant rs705704.
PP = posterior probability of colocalisation between the QTL (eQTL or caQTL) and the T1D association (referred to in *coloc* documentation as “PP.H4.abf”)
caQTL = chromatin accessibility quantitative trait locus
eQTL = expression quantitative trait locus

### Functional annotation of TID-associated variants in the *BACH2* region

Fine-mapping of the *BACH2* locus refined the T1D association to two intronic variants, rs72928038 (G>A) and rs6908626 (G>T) (**Figure 3a**). The EUR minor alleles of rs72928038 (A) and rs6908626 (T) are associated with increased T1D risk (OR = 1.18; p < 1×10^-20^, MAF_EUR_ = 0.18). Chromatin-state annotations across cell types from the BLUEPRINT Consortium and NIH Roadmap Epigenomics Project annotate rs72928038 as overlapping a T cell-specific active enhancer and rs6908626 as lying in the ubiquitous *BACH2* promoter (**Figure 3b**, https://github.com/ccrobertson/t1d-immunochip-2020 - see “BACH2”). Promoter-capture Hi-C data from diverse immune cell types^34^ indicates that the enhancer region containing rs72928038 contacts the *BACH2* promoter in T cells (**Figure 3c**). Although weak interactions were observed in multiple T cell subtypes, only naïve CD4^+^ T cells had a significant interaction score.

In caQTL analysis, rs72928038 (A) is associated with decreased accessibility of the enhancer it overlaps (chr6:90266766-90267715) (**Figure 3d - left**), while rs6908626 (T) does not appear to affect accessibility at the *BACH2* promoter (chr6:90294665-90297341) (**Figure 3d – right**). Similarly, among 15 samples heterozygous for rs72928038, only 4% (5/121) of ATAC-seq reads overlapping that site contain the T1D risk allele, rs72928038 (A) (**Figure 3e – left, Supplementary Table 15**), suggesting it leads to restricted accessibility. In contrast, chromatin accessibility at rs6908626 does not exhibit allelic bias in heterozygotes (**Figure 3e – right**). These data prioritise rs72928038 over rs6908626 as functionally relevant in CD4^+^ T cells.

**Figure 3:**
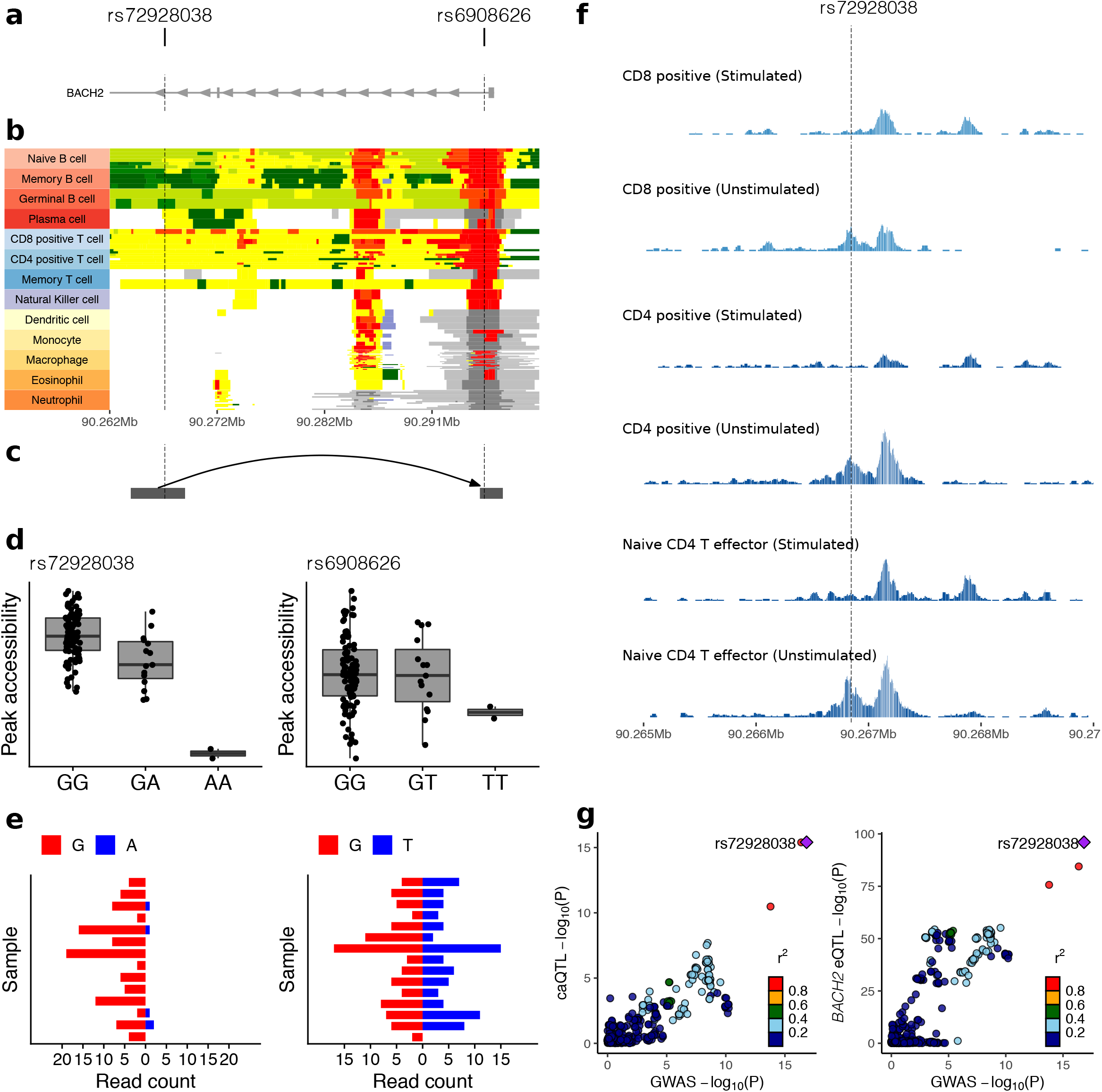
Functional annotation of T1D-associated variants in the *BACH2* region. **a)** T1D-associated variants prioritised by fine-mapping (rs72928038 and rs6908262) and overlap with introns of *BACH2*. **b)** chromHMM tracks across diverse immune cell types from the BLUEPRINT consortium; rs72928038 resides in a T cell-specific enhancer (yellow regions), while rs6908262 overlaps the *BACH2* promoter (red regions). **c)** Interactions with the *BACH2* promoter in published PCHi-C data from naïve CD4^+^ T cells^34^. **d)** Accessibility of regions overlapping rs72928038 and rs6908262 by genotype. The T1D risk alleles, rs72928038(A) and rs6908262(T), are associated with decreased chromatin accessibility of the region overlapping rs72928038 (left), but not with accessibility in the region overlapping rs6908262 (right). **e)** Allele-specific accessibility of chromatin at rs72928038 and rs6908262 across heterozygous individuals; at rs72928038, the G allele is strongly preferred over the A allele. **f)** Chromatin accessibility profiles in the region overlapping rs72928038 across resting and activated CD4^+^ and CD8^+^ T cells (published data^29^); chr6:90266766-90267715 is accessible specifically in unstimulated cells. **g)** LocusCompare plots showing colocalization between T1D association, the caQTL for chr6:90266766-90267715 (left), and the eQTL for *BACH2* (right).

In eQTL studies, rs72928038 (A) is associated with decreased expression of *BACH2* in whole blood^33^ and purified immune cell types^35^. Specifically, in the DICE consortium^35^, rs72928038 (A) is associated with decreased expression of *BACH2* in multiple cell types, with the strongest effects in naïve CD4^+^ and CD8^+^ T cells. This is consistent with the observation that the enhancer region overlapping rs72928038 is accessible specifically in unstimulated bulk CD4^+^, unstimulated bulk CD8^+^, and naïve CD4^+^ T effector cells (**Figure 3f**). Both the enhancer caQTL and *BACH2* eQTL colocalize with T1D association (**Figure 3g, Table 2**).

The *BACH2* rs72928038 variant overlaps binding sites for STAT1 and the ETS family of transcription factors, based on canonical transcription factor binding motifs^25^. We performed super-shift electrophoretic mobility shift assay (EMSA) experiments of the DNA sequence flanking rs72928038 that demonstrated allele-specific ETS1 binding, but no STAT1 binding (**Supplementary Figure 10**). This result builds on experiments demonstrating allele-specific nuclear protein binding of rs72928038 in Jurkat cells^36^. These data prioritise rs72928038 as the functionally relevant variant in T cells and provide preliminary support for a gene regulatory mechanism underlying the 6q15 region association with T1D where the rs72928038 minor A allele disrupts ETS1 binding, which leads to decreased enhancer activity and *BACH2* expression in naïve CD4^+^ T cells.

### T1D drug target identification

To identify potential T1D therapeutic targets with human genetic support, we used the Priority Index (Pi) algorithm,^17^ which integrates genetic association results with functional genomics data and protein-protein networks (Online Methods). We identified 50 highly-ranked gene targets (**Supplementary Table 16**), including 13 that were not previously implicated by Priority Index analyses^17^: *STAT4, RGS1, CXCR6, IL23A, PTPN22, NFKB1, MAPK3, EPOR, DGKQ, GALT, IL12RB1, IL12RB2, IL6R*, and 12 that have been targeted in clinical trials for autoimmune diseases: *IL2RA, IL6ST, IL6R, TYK2, IFNAR2, JAK2, IL12B, IL23A, IL2RG, JAK3, JAK1* and *IL2RB*. These results provide genetic support for exploring these genes and pathways as potential therapeutic targets for T1D prevention or treatment.

## Discussion

In the largest genetic analysis of T1D to date, we identified 36 novel regions at p < 5×10^-8^ and implicated a total of 152 regions outside the MHC in T1D susceptibility at FDR < 0.01. Additionally, we refined the set of putative causal variants and number of independent associations in many T1D regions through increased sample size, dense genotyping and imputation, inclusion of multiple ancestry groups, and optimized analytical approaches to fine-mapping. We assessed their intersection with regions of putative regulatory function with public and newly generated ATAC-seq data from diverse cell types and states. T1D credible variants were enriched in stimulation-responsive open chromatin peaks in CD4^+^ T cells. Colocalization of T1D associations with CD4^+^ T cell caQTLs focused mechanistic hypotheses on this highly relevant cell type. Four of the five T1D associations that colocalize with caQTLs also colocalize with whole-blood eQTLs. These colocalized associations offer hypotheses of how causal variants influence disease risk through their effects on regulatory element activity and gene expression in T1D-relevent cell types. We propose the regulatory mechanism by which the disease association in the *BACH2* locus leads to disease risk. Finally, using our genetic association data coupled with functional genomics and protein networks, we identified potential T1D drug targets for use in prevention trials. Experimental follow-up studies are required to test these hypotheses and further dissect the mechanisms altering T1D risk in each region.

We highlight the *BACH2* region on chromosome 6q15 as an example of unbiased QTL colocalization that leads to hypotheses for functional mechanisms driving variant-T1D association. Our data suggest that rs72928038 (A), the T1D-associated allele, abolishes ETS1 binding at an enhancer that promotes *BACH2* expression in naïve CD4^+^ T cells. *BACH2* is a transcription factor from the BTB-basic leucine zipper family with established roles in B and T cell biology, including maintaining the naïve T cell state^37,38^. *BACH2* haploinsufficiency has been shown to cause congenital autoimmunity and immunodeficiency^39^, demonstrating that a functioning human immune system depends on *BACH2* expression in a dose-dependent manner. In addition to *cis*-effects on *BACH2* expression, rs72928038 is associated with changes in expression of 39 distal genes^33^ in whole blood, including seven genes in autoimmune disease-associated regions. These observations raise the hypothesis that the minor A allele at rs72928038 increases T1D (and other autoimmune disease) risk by reducing *BACH2* expression in a precise cellular context (e.g., the naïve T cell state). This effect may lead to shifts in *BACH2*-regulated transcriptional programs, thereby altering T cell lineage differentiation in response to antigen exposure.

Unsurprisingly, given the design of the ImmunoChip, 13 of the 36 novel (p < 5×10^-8^) regions associated with T1D risk are associated with other immune-related traits. This is consistent with previous studies showing shared genetic risk across autoimmune disease^40^ and suggests potential for repurposing drugs to treat or prevent T1D. This was further highlighted by the results from our Priority Index target prioritisation analysis, where 12 targets were prioritised that have been the focus of clinical trials for treatment of autoimmune diseases. One example is *IL23A*, which has been targeted in the treatment of irritable bowel disease (IBD), psoriasis, multiple sclerosis and ankylosing spondylitis. The IL-23 inhibitors have been efficacious and safe in the treatment of IBD^41^ and psoriasis^42^, and are currently being explored for T1D (ClinicalTrials.gov identifiers NCT02204397 and NCT03941132); the present analysis gives genetic support for these trials. Similarly, *JAK1, JAK2* and *JAK3* were implicated in T1D etiology in our analysis. JAK inhibitors have been safe and effective in the treatment of rheumatoid arthritis^43^ and ulcerative colitis^44^. Finally, this study presents the first well-powered, convincing genetic evidence linking interleukin-6 (IL-6), a cytokine with known roles in multiple autoimmune diseases, to T1D etiology. The IL-6 receptor complex consists of two essential subunits: the alpha subunit (encoded by *IL6R)* and the signal transducing subunit (encoded by *IL6ST*). Both the *IL6ST* and *IL6R* regions were identified here as T1D-associated at genome-wide significance for the first time (Table 1). Additionally, both *IL6ST* and *IL6R* were implicated by our Priority Index analysis. We cannot say based on current evidence that *IL6ST* and *IL6R* are T1D causal genes, but we note that *IL6ST* is implicated by QTL colocalisation and the lead T1D variant near *IL6R* (rs2229238) is an eQTL for *IL6R* expression in whole blood (formal colocalisation was not assessed because the *IL6R* region is not densely covered by the ImmunoChip). The humanized IL-6 receptor antagonist monoclonal antibody, tocilizumab, is an approved treatment for RA and systemic juvenile idiopathic arthritis, both of which share substantial genetic effects with T1D^3^, and a trial of this drug in recently-diagnosed T1D cases is underway (https://clinicaltrials.gov/ct2/show/NCT02293837). However, surprisingly we showed that the lead T1D variant near *IL6R* (rs2229238) tags a causal variant distinct from the nonsynonymous variant (*IL6R* Asp358Ala; rs2228145 A>C) thought to drive the association in RA^19^, suggesting potentially different mechanisms altering disease risk in this region. The recent success of anti-CD3 therapy, after 40 years of study through experimental models and clinical trials targeting different patient subgroups and time points relative to disease diagnosis^45^, highlights both the challenges and hopes for translating target identification to efficacious clinical outcomes in T1D.

One limitation on this study is that genotyping was restricted to ImmunoChip content, which provides dense coverage in 188 immune-relevant genomic regions, as defined by previous largely European-based GWAS of immune-related traits, and sparse coverage elsewhere. This design restricts the scope of discovery, fine-mapping, and generalizability of subsequent functional enrichment analyses. This may explain the absence of T1D variant enrichment in open chromatin of non-immune cell types (e.g., pancreatic islets)^46,47^. While this analysis is the largest and most comprehensive study prioritising novel therapeutic interventions in T1D according to genetic evidence, extension of future genetic studies to genome-wide analyses and continuing to focus on more diverse populations will further define the genetic landscape of T1D.

## Author contributions

The study was conceptually designed by P.C., J.A.T. and S.S.R.

The study was implemented by S.O.G., P.C., J.A.T. and S.S.R.

DNA samples for genotyping were managed by S.O.G and E.F.

Frozen T1DGC PBMC samples for chromatin accessibility profiling (ATAC-seq) were managed by P.C. and S.O.G.

ATAC-seq data generation at University of Virginia was led by S.O.G.

ATAC-seq data generation at University of Oxford was led by A.J.C.

Genotype data processing, quality control, imputation, and statistical analyses were performed by W-M.C., S.O.G., J.R.J.I. and C.C.R.

Chromatin accessibility data processing and analysis was performed by A.J.C., D.F.S.C., J.R.J.I., and C.C.R.

EMSA assays were performed by H.Y., with supervision from S.O.G.

D.B.D. provided samples for genotyping through the Genetic Resource Investigating Diabetes (GRID).

P.D. provided ImmunoChip genotyping data through the UK Blood Service (UKBS).

J.H.B. provided samples for genotyping and data from the Benaroya Research Institute (BRI).

S.L.B. provided samples for genotyping through the Consortium for the Longitudinal Evaluation of African-Americans with Early Rheumatoid Arthritis (CLEAR).

P.K.G. provided samples for genotyping through the New York Cancer Project (NYCP).

J.D., D.D., J.M.L., S.M., and A.S.S. provided samples for genotyping and data through the SEARCH for Diabetes in Youth study (SEARCH).

C.J.G. and M.A.A. provided samples for genotyping through the Type 1 Diabetes TrialNet study (TrialNet).

R.P.K., J.C.E., M.J.R., A.K.S., J.R.O., and F.P. provided samples for genotyping through their affiliated institutions and research programs.

The manuscript was written by J.R.J.I. (under supervision by J.A.T.) and C.C.R. (under supervision by S.S.R.).

All authors reviewed and approved the manuscript.

## Acknowledgements

We thank the investigators and their studies for contributing samples and/or data to the current work, and the participants in those studies who made this research possible. These studies include T1DGC, British 1958 Birth Cohort, GRID, CLEAR, Epidemiology of Diabetes Interventions and Complications (EDIC), Genetics of Kidneys and Diabetes Study (GoKinD), NYCP, SEARCH, TrialNet, Tyypin 1 Diabetekseen Sairastuneita Perheenjäsenineen (IDDMGEN), Tyypin 1 Diabeteksen Genetiikka (T1DGEN), Northern Ireland GRID Collection, Northern Ireland Young Hearts Project, Hvidoere Study Group on Childhood Diabetes (HSG), and International HapMap Project. Additional institutions contributing samples are: British Diabetes Association (BDA), NIHR Cambridge BioResource, UKBS, BRI, National Institute of Mental Health (NIMH), University of Alabama at Birmingham (UAB), University of Colorado (UC), University of California San Francisco (UCSF), Medical College of Wisconsin (MCW), and Steno Diabetes Center. Samples and data can be obtained on T1DGC, EDIC, and GoKinD from the NIDDK Central Repository.

This research utilizes resources provided by the T1DGC, a collaborative clinical study sponsored by the National Institute of Diabetes and Digestive and Kidney Diseases (NIDDK), National Institute of Allergy and Infectious Diseases, National Human Genome Research Institute, National Institute of Child Health and Human Development, and JDRF and supported by U01 DK-062418. The generation of chromatin accessibility data on T1DGC samples was supported by grants from the NIDDK (DP3-111906 to S.S.R. and DK-115694 to P.C.).

While working on this project, C.C.R was supported by a training grant from the U.S. National Library of Medicine (5T32LM012416) and the Wagner Fellowship from the University of Virginia.

This work made use of data and samples generated by the 1958 Birth Cohort (NCDS), which is managed by the Centre for Longitudinal Studies at the UCL Institute of Education, funded by the Economic and Social Research Council (grant number ES/M001660/1). Access to these resources was enabled via the Wellcome Trust & MRC: 58FORWARDS grant [108439/Z/15/Z] (The 1958 Birth Cohort: Fostering new Opportunities for Research via Wider Access to Reliable Data and Samples). Before 2015 biomedical resources were maintained under the Wellcome Trust and Medical Research Council 58READIE Project (grant numbers WT095219MA and G1001799).

We acknowledge use of DNA samples from the NIHR Cambridge BioResource. We thank volunteers for their support and participation in the Cambridge BioResource and members of the Cambridge BioResource Scientific Advisory Board (SAB) and Management Committee for their support of our study. We acknowledge the NIHR Cambridge Biomedical Research Centre for funding. Access to Cambridge BioResource volunteers and to their data and samples are governed by the Cambridge BioResource SAB. Documents describing access arrangements and contact details are available at http://www.cambridgebioresource.org.uk/.

The ethics for GRID were processed by the NRES Committee East Of England Cambridge South MREC 00/5/44.

The authors thank the following CLEAR investigators who performed recruiting: Drs. Doyt Conn (Grady Hospital and Emory University, Atlanta, GA), Beth Jonas and Leigh Callahan (University of North Carolina at Chapel Hill, Chapel Hill, NC), Edwin Smith (Medical University of South Carolina, Charleston, SC), Richard Brasington (Washington University, St. Louis, MO), and Larry W. Moreland (University of Pittsburgh). The CLEAR Registry and Repository was funded by National Institutes of Health (NIH) Office of the Director grants N01-AR-0-2247 (9/30/2000–9/29/2006) and N01 AR-6-2278 (9/30/2006–3/31/2012) (S.L.B., PI).

Bio-samples and/or data for this publication were obtained from NIMH Repository & Genomics Resource, a centralized national biorepository for genetic studies of psychiatric disorders.

The SEARCH for Diabetes in Youth Study (www.searchfordiabetes.org) is indebted to the many youth and their families, as well as their health care providers, whose participation made this study possible. SEARCH for Diabetes in Youth is funded by the Centers for Disease Control and Prevention (PA numbers 00097, DP-05-069, and DP-10-001) and supported by the NIDDK. SEARCH Site Contract Numbers: Kaiser Permanente Southern California (U48/CCU919219, U01 DP000246, and U18DP002714), University of Colorado Denver (U48/CCU819241-3, U01 DP000247, and U18DP000247-06A1), Children’s Hospital Medical Center (Cincinnati) (U48/CCU519239, U01 DP000248, and 1U18DP002709), University of North Carolina at Chapel Hill (U48/CCU419249, U01 DP000254, and U18DP002708), University of Washington School of Medicine (U58/CCU019235-4, U01 DP000244, and U18DP002710-01), Wake Forest University School of Medicine (U48/CCU919219, U01 DP000250, and 200-2010-35171).

We acknowledge the support of the Type 1 Diabetes TrialNet Study Group (https://www.trialnet.org), which identified study participants and provided samples and followup data for this study. The Type 1 Diabetes TrialNet Study Group is a clinical trials network funded by the National Institutes of Health (NIH) through the National Institute of Diabetes and Digestive and Kidney Diseases, the National Institute of Allergy and Infectious Diseases, and The Eunice Kennedy Shriver National Institute of Child Health and Human Development, through the cooperative agreements U01 DK061010, U01 DK061016, U01 DK061034, U01 DK061036, U01 DK061040, U01 DK061041, U01 DK061042, U01 DK061055, U01 DK061058, U01 DK084565, U01 DK085453, U01 DK085461, U01 DK085463, U01 DK085466, U01 DK085499, U01 DK085505, U01 DK085509, and JDRF. The contents of this article are solely the responsibility of the authors and do not necessarily represent the official views of the NIH or JDRF.

DNA samples from UAB were recruited, in part, with the support of P01-AR49084 (R.P.K., principal investigator [PI]) and of UL1-TR001417 (R.P.K., PI).

We acknowledge the involvement of University of Colorado Pediatric Clinical and Translational Research Center (NIH/NCATS grant number UL1 TR000154), the Barbara Davis Center at the University of Colorado Denver (DERC NIH grant number P30 DK57516).

WGS data production and variant calling was funded by an NHGRI Center for Common Disease Genomics award to Washington University in St. Louis (UM1 HG008853).

This study used the Trans-Omics in Precision Medicine (TOPMed) program imputation panel (version TOPMed-r2) supported by the National Heart, Lung and Blood Institute (NHLBI); see www.nhlbiwgs.org. TOPMed study investigators contributed data to the reference panel, which can be accessed through the Michigan Imputation Server; see https://imputationserver.sph.umich.edu. The panel was constructed and implemented by the TOPMed Informatics Research Center at the University of Michigan (3R01HL-117626-02S1; contract HHSN268201800002I). The TOPMed Data Coordinating Center (3R01HL-120393-02S1; contract HHSN268201800001I) provided additional data management, sample identity checks, and overall program coordination and support. We gratefully acknowledge the studies and participants who provided biological samples and data for TOPMed.

