## Supplementary material for "Fine-mapping, trans-ancestral and genomic analyses identify causal variants, cells, genes and drug targets for type 1 diabetes": Online Methods

### Genotyping and quality control

DNA samples were genotyped on the Illumina ImmunoChip at University of Virginia Genome Sciences Laboratory (N = 52,219), Sanger Institute (N = 4,347), University of Cambridge (N = 2,941), and Feinstein Institute (N = 1,811). Raw genotyping files were assembled at the University of Virginia. Genotype clusters were generated using the Illumina GeneTrain2 algorithm at University of Virginia. Stringent SNP- and sample-level quality control filtering and data cleaning was performed to ensure high quality genotypes and accurate pedigrees (**Supplementary Figure 11**). Specifically, we applied the following variant filters:

(1) re-annotated ImmunoChip variant positions by aligning probe sequences to hg37 and removed any variants with less than a 100% match or multiple matches at different positions in the genome;

(2) removed variants with call rates less than 98%;

(3) removed variants with any discordance between duplicate or monozygotic twin samples, as confirmed by genotype-inferred relationships;

(4) removed variants with Mendelian inconsistencies in more than 1% of informative trios or parent-offspring pairs, based on genotype-inferred relationships.

For sample filtering, we used X chromosome heterozygosity and Y chromosome missingness to identify and exclude participants with apparent sex chromosome anomalies or resolve inconsistencies with reported sex. Additionally, pedigree-defined and genotype-inferred sample relationships were compared using the software KING[1] (http://people.virginia.edu/~wc9c/KING) and samples were excluded when inconsistencies could not be resolved. Relationships between families, within and across cohorts, were also checked. For each pair of related families observed, we randomly selected one to remove from association analysis. After resolving sex and relationship issues, samples with genotype call rate less than 98% were removed and variants with genotype frequencies deviating from Hardy-Weinberg Equilibrium ($p < 5\times10^{-5}$) in unrelated European controls were excluded before imputation.

**Ancestry analyses**

Principal components (PC) were generated in 1000 Genomes phase 3 individuals using 8,297 autosomal ImmunoChip variants selected by excluding regions of long-range linkage disequilibrium (LD) [2], pruning for short-range LD ($R^2 < 0.2$ in 50kb windows), and filtering for MAF (> 0.05). The participants were then projected onto the 1000 Genomes PC space using PLINK v1.9 [3]. The first ten PCs were then used in k-means clustering to define five clusters of ancestrally-similar participants, namely European (EUR), African-American (AFR), East Asian (EAS), Finnish (FIN) and Admixed (AMR), labeled according to their closest 1000 Genomes super-population.

**Whole genome sequencing**

Whole genome sequencing (WGS) data were available in a subset of samples, including 1,411 AFR, 641 AMR, and 95 EUR subjects, through the NHGRI Centers for Common Disease Genomics (CCDG) (https://ccdg.rutgers.edu/). Samples were sequenced on the Illumina HiSeq X at the McDonnell Genome Institute at Washington University in St. Louis. Sequence alignment and variant calling was performed as outlined in the standardized CCDG pipeline [4] (https://github.com/CCDG/Pipeline-Standardization/blob/master/PipelineStandard.md).

**Identification and stratification of unrelated individuals and family trios**

Case-control analysis was performed separately for each ancestry cluster. Within each cluster, affected trios were excluded and a set of unrelated individuals was selected from the remaining subjects using KING software ("--unrelated" option) [1]. Cluster-specific PCs were calculated by performing PC analysis on unrelated controls and projecting the remaining subjects onto the resulting axes. Remaining population stratification within each ancestry cluster was assessed visually (**Supplementary Figure 12**).

**Imputation to TOPMed and 1000 Genomes reference panels**

Genotypes were imputed across the entirety of all autosomal chromosomes, with the NHLBI Trans-Omics for Precision Medicine (TOPMed) Freeze 5 [5] and 1000 Genomes phase 3 reference panels using the Michigan Imputation Server, which applied Eagle version 2.4 [6] for phasing and Minimac4 for imputation[7]. For each reference panel, ImmunoChip variants were aligned to the appropriate strands and reference alleles using available tools (https://www.well.ox.ac.uk/~wrayner/tools/). Since imputation quality and R-squared statistics are dependent on allele frequency and LD patterns in the target population, we filtered imputed variants for ancestry-specific imputation quality ($R^2 > 0.8$ and INFO > 0.8), MAF (> 0.005), and Mendelian inconsistency rates (< 0.01 in informative trios and parent-offspring pairs, only considered in the family-based association analyses).

**Defining ImmunoChip regions**

The ImmunoChip was designed to densely cover genetic variation in approximately 188 immune-associated genomic regions. We mapped 188 previously defined “ImmunoChip regions” (defined from the “humarray” R package) from GRCh36 to GRCh38 coordinates (**Supplementary Table 2**): for each region, we mapped all variants originally included in the region to GRCh38 and then defined the boundaries of the region as the lowest and highest observed GRCh38 positions among these variants (+/- 50 kb either side).

We performed both association discovery analyses and fine-mapping analyses in ImmunoChip regions.

**Defining other densely genotyped regions**

We defined other densely genotype regions as any 500kb region on the ImmunoChip that contained more than 50 variants (**Supplementary Table 3**). Although these regions were not genotyped at the density observed in ImmunoChip regions, we considered them to be contributing enough information to the imputation to obtain reliable association summary statistics from variants imputed in these regions.

We performed association discovery analyses in densely genotyped regions, including imputed variants passing imputation quality filters, but did not attempt to fine-map these regions (unless they overlapped an ImmunoChip region).

**Association analysis – Phase I**

We analyzed association of each variant with type 1 diabetes (T1D) separately in the five ancestry groups, assuming an additive mode of inheritance and using logistic regression for unrelated case-control analyses, adjusting for five ancestry-specific PCs and using genotype posterior probabilities to account for uncertainty in imputed genotypes using the SNPTEST software [8]. Individuals genetically clustering into the EAS ancestry group were excluded from this analysis, since there were too few individuals in the analysis to obtain stable estimates (38 cases and 106 controls). We combined results from each ancestry in an inverse-variance weighted fixed-effects meta-analysis, using the METAL software [9].

A number of variant quality control measures were applied to prevent spurious associations due to imputation: variants with MAF < 0.005 were excluded, as well as variants with a SNPTEST info score < 0.8 [8] (in cases, controls or overall). Variants with a difference in SNPTEST info score > 0.05 between cases and controls were removed since this could artificially generate an association that is reflecting imputation differences rather than genuine differences in allele frequencies between cases and controls. Finally, only imputed variants lying within the 188 "ImmunoChip regions" (**Supplementary Table 2**) or in other densely genotyped regions outside of the ImmunoChip regions defined above (**Supplementary Table 3**) were analyzed for association with T1D, since genotyping outside these regions on the ImmunoChip is sparse and therefore imputed variant calls less certain. However, all variants that were directly genotyped and passed QC on the ImmunoChip were included in the association analysis.

Forward stepwise logistic regression was performed to identify loci with more than one independent association with T1D. All conditionally independent associations ($p < 5\times10^{-8}$) were reported. Case-control analyses were performed under recessive, and dominant models of inheritance. To evaluate the relative fit of the three models, we compared the Akaike information criterion (AIC) in the EUR ancestry group and identified the model providing the lowest AIC, which indicates the best fit.

On the X chromosome, only genotyped variants were examined for their association with T1D, and the Y chromosome was not examined.

**Association analysis – trio families and combined analyses**

The trio families (two parents and an affected offspring) were analyzed by ancestry group, and the transmission disequilibrium test (TDT) [10] was used to test for association between each variant and T1D. As previously shown, TDT statistics are susceptible to substantial bias when applied to imputed genotypes [11]. To mitigate this, we applied a stringent variant filter to the imputed genotypes, removing all variants with Mendelian inconsistencies in > 1% of trios with heterozygous offspring or parent-offspring pairs with homozygous offspring. From TDT summary statistics, we derived effect sizes and standard error estimates for each variant to enable meta-analysis with the additive model, case-control meta-analysis results, as previously demonstrated [12].

**Statistical fine-mapping**

We used two complementary approaches to define credible variant sets within each T1D-associated ImmunoChip region. To define the fine-mapping region to examine, we took the lead variant and examined all variants that were included in the discovery analysis within 1.5Mb of the lead variant (750kb either side). This would usually consist of the entirety of the ImmunoChip region and also any other variants directly genotyped on the ImmunoChip that were proximal to the ImmunoChip region.

*1) Fine-mapping using European Case-Control data only (GUESSFM)*

Since forward stepwise model selection can fail to identify complex genetic architectures [13], we applied a Bayesian method (GUESSFM) in the EUR case-control data, which attempts to identify the most likely combinations of variants explaining the phenotype [14,15].

Briefly, the ($n \times m$) genetic matrix $\mathbf{X}$ at a locus, where $n$ is the number of individuals and $m$ is the number of variants, is pruned to remove variants in high LD ($r^2 \geq 0.99$), generating a pruned ($n \times p$) matrix, $\mathbf{Z}$, with $p$ 'tag' variants. Now the model space contains $2^p$ possible models. A stochastic search is carried out across these $2^p$ models, averaging over the other parameters (e.g. variant effect size), to obtain a selection of models with high marginal posterior probabilities. This set of models is then expanded to include models where the tag variant is replaced by each of the variants in high LD with it (which had been removed during pruning prior to the stochastic search). For each model, an Approximate Bayes Factor (ABF) is calculated by treating the binary outcome (T1D status) as linear and using the linear regression Bayesian Information Criterion. The ABF for each model can be interpreted as the support for the model relative to a null model with no genetic variants. To obtain the posterior probability for each model, the ABF is multiplied by the prior, which we took in all cases to be a binomial prior with $3/m$ expected variants included in the model, divided by the normalizing factor, the sum of all tested model posterior probabilities. The marginal probability for each SNP is taken as the sum of the posterior probabilities for all models in which it is present.

In the results, we refer to groups of variants prioritised by GUESSFM as "credible sets" and variants within these groups as "credible variants." Variants that failed quality control metrics (or were not genotyped or imputed in our data for other reasons) but are in LD ($r^2 > 0.9$ in 1000

Genomes Phase 3) with a prioritised variant are also included in the comprehensive list of credible variants provided in **Supplementary Table 11**.

*2) Trans-ethnic fine-mapping*

In regions where association signals were marginally associated ($p < 5\times10^{-4}$) in multiple ancestry groups and where the evidence from EUR ancestry only fine-mapping suggested a single causal variant in the region (marginal posterior probability for one causal variant in the region > 0.5), we applied the multi-ethnic fine-mapping method, PAINTOR [16] to further refine association signals. PAINTOR uses association z-scores and population level LD estimates to identify the combination of alleles that best explain the phenotype, multiplying the posterior probability of the casual vector across ancestry groups, assuming the same variant(s) are causal in each ancestry group. Since loci examined were those with evidence of one causal variant in the region, we restricted the maximum model size to two variants in the region and enumerated the posterior of every model rather than performing an MCMC search. The association z-scores used for each ancestry group were from a meta-analysis of case-controls and family trios in that ancestry cluster.

**Haplotype analyses**

Haplotype analyses were performed by taking "best-guess" genotype values for the variants included in the analysis and obtaining haplotype phase distribution estimates for each individual, using an expectation-maximization algorithm [17]. Each individual's haplotype was then sampled ten times and a logistic regression was fitted estimating the effect size of the haplotype relative to the most common haplotype in the population, with T1D status as the outcome and adjusting for the five largest genetic PCs. The estimates and standard errors for each haplotype relative to the most common were then averaged over these ten logistic regression models to obtain overall haplotype effect sizes on T1D risk. These analyses were performed using the EUR ancestry cases and controls only.

**Annotating T1D-associated protein-altering variants**

For each variant with group posterior probability > 0.5 in the comprehensive list of T1D credible variants (**Supplementary Table 11**), we annotated the functional impact using the software ANNOVAR [18] and the Ensembl and refGene annotation databases.

**Obtaining publicly available data**

*ATAC-seq.*

Raw FASTQ files were obtained from Gene Expression Omnibus (GEO) accession number GSE118189. These data included four individuals and 25 immune cell types under resting conditions and after stimulation with anti-human CD3/CD28 dynabeads and human IL-2 (for 24 hours, T lymphocytes), F(ab)'2 anti-human IgG/IgM [19] and human IL-4 (for 24 hours, B lymphocytes), human IL-2 (for 48 hours, NK cells), or LPS (for 6 hours, monocytes)[20].

ATAC-seq data from pancreatic islets of five donors without glucose intolerance and five EndoCβH1 cell line replicates, under resting conditions and after stimulation with IFN-γ and IL-1β for 48 hours were downloaded from GEO, accession number GSE123404 [21].

ATAC-seq data from cardiac fibroblasts (two fetal and three adult) were downloaded from the European Nucleotide Archive (https://www.ebi.ac.uk/ena/data/view/SRX2843570 and https://www.ebi.ac.uk/ena/data/view/SRX2843571), as a control cell type that we did not expect to be involved in the aetiology of T1D [22].

*Epigenome annotation tracks.*

chromHMM [23] tracks from diverse primary human cells were obtained from the NIH Epigenome Roadmap, http://dcc.blueprint-epigenome.eu/#/md/secondary_analysis/Segmentation_of_ChIP-Seq_data_20140811

and additional immune-specific human primary and cell lines from the Blueprint consortium, https://egg2.wustl.edu/roadmap/data/byFileType/chromhmmSegmentations/ChmmModels/coreMarks/jointModel/final.

*Whole blood eQTL summary statistic*s.

Summary statistics from whole blood *cis* eQTL analysis from 31,683 individuals[24] were downloaded from https://eqtlgen.org.

**Generating additional ATAC-seq data for enrichment analyses**

In addition to using publicly-available ATAC-seq data to examine T1D variant enrichment [19], ATAC-seq data were also generated in Oxford, using different culture and stimulation conditions. Two cell types were examined; $CD4^+$ T cells (n=6) and $CD19^+$ B cells (n=4). $CD4^+$ T cells were enriched and stimulated as previously described [25]. B cells were positively selected from PBMCs using anti-CD19 beads (Miltenyi Biotec, GmbH) and cultured for 24 hours in X-VIVO 15 (Lonza, Switzerland) supplemented with 1% Human Ab Serum (Sigma) and penicillin / streptomycin (Thermo Fisher) and plated in 96-well CELLSTAR U–bottomed plates (Greiner

Bio-One, Austria) at concentration of 2.5×105 cells/well. Cells were left untreated or stimulated with goat anti-human IgM/IgG/IgA antibody (10ug/ml, ENZO lifesciences), rhIL-21 and rhIL-4 (20ng/ml, Peprotech) for 24 hours. ATAC-seq data was generated from 50,000 cells from each cell type and culture condition following the Omni-ATAC protocol [26].

**Processing ATAC-seq data for enrichment analyses**

ATAC-seq datasets were mapped with minimap2 (version 2.17) [27], except for GSE123404 (pancreatic islets dataset) where bowtie2 (version 2.3.5) was used. In all cases, the mapping was performed against the reference genome GRCh38.p12, obtained from the GENCODE project [28]. After mapping, the technical replicates (where available) were merged with Picard tools (version 2.20.2). PCR duplicated reads were detected and removed with the same tool. The percentage of detected duplicated reads was very low in all datasets, with a mean value < 1%.

For visualization purposes, a set of bigwig files were generated with bamCoverage from the deeptools package (version 3.3.0), using reads per genome coverage (RPGC) normalization and ignoring allosomes and the mitochondrial chromosome. Peaks were called using macs2 [29] (version 2.1.2) with the params "--nomodel --shift 37 --extsize 73 --keep-dup all".

The immune cell ATAC-seq dataset GSE118189 [19] was used to create a consensus list of peaks. For each cell type, we selected the donor contributing the fewest number of reads to that cell type and divided this number by two. Reads were then randomly pooled by that number for each sample, creating a representative alignment file for that cell type. This procedure was performed twice in order to obtain two pseudo-replicates. Peaks were called with macs2 with the same parameters. Irreducible discovery rate (IDR) was calculated between the two pseudo replicates [30], any peak with an IDR ≤ 0.05 were included in the consensus list of peaks. This list was then used as a feature reference and reads were counted per feature with featureCounts from the package subread [31] (version 1.6.4). A similar approach was used for the other datasets in the analysis. IDR was used to obtain a reliable list of peaks. In these datasets, no feature reference was derived from the IDR, and counting was performed directly from the list obtained from GSE118189.

To orchestrate these workflows, conda and snakemake were used. The code with the conda packages needed to run the workflow have been uploaded to https://github.com/dfloresDIL/MEGA.

**ATAC-seq enrichment analyses**

To examine enrichment of T1D credible variants (group marginal posterior probability > 0.8 from GUESSFM) in open chromatin, for each cell type we compared the number of variants falling within open chromatin among T1D credible variants versus variants in regions of the genome with similar LD structure and gene density. To do this, we did the following:

1) Using European individuals from 1000 Genomes Project data, identified all variants with a Pearson correlation > 0.8 with each other.
2) Binned the T1D credible variants with group marginal posterior probability > 0.8 with regards to their LD block size: 1-9, 10-19, 20-49, 50-74,75-99,100-149 or 150-249.
3) Binned the 1000 Genomes Project data variants in the same way with regards to LD block size, taking an LD block as the variants with Pearson correlation > 0.8 with an index variant.

4) For each T1D credible group, selected randomly an LD block from the 1000 Genomes Project data of the same bin size and with the same (or similar for large haplotypes) number of genes overlapping the credible group, therefore selecting a similar number of variants to the T1D credible group, with an approximately equivalent LD structure and gene density.

5) Repeated step 4) 100 times, therefore selecting 100 randomly sampled parts of the genome with approximately equivalent size and LD structure to the T1D credible variants.

6) For cell type, X, counted the number of T1D credible SNPs overlapping ATAC-seq peaks. Compared this to the number overlapping ATAC-seq peaks from the first randomly sampled set of variants. Performed a Fisher's exact test comparing the intersection with ATAC-seq peaks between the T1D credible variants and the randomly sampled variants with equivalent size, gene density and LD structure and obtained a z-score from this test.

7) Repeated step 6) 100 times, one for each randomly sampled set of haplotypes across the genome, obtaining 100 z-scores.

8) Took the mean z-score from the 100 tests and compared it to a normal distribution to get an overall enrichment p-value for cell type X.

We performed steps 6) to 8) for each cell type and under each condition.

*Condition specific enrichment analyses*. Since many ATAC-seq peaks are present in stimulated and unstimulated conditions, the enrichment scores are similar across conditions. However, there are peaks that are differentially-open between conditions, which the initial enrichment analysis does not account for. Therefore, for cell types with data available from unstimulated and stimulated conditions, we tested for differential-expression of counts between the conditions using DESeq2 [32], examining only peaks in the consensus list of peaks from ATAC-seq dataset

GSE118189 (25 immune cell types). Those peaks with association FDR<0.01 were examined for their enrichment for T1D credible variants using steps 6) to 8) above.

**Generating ATAC-seq data in patient samples for caQTL analyses**

To assess whether T1D-associated variants affect chromatin accessibility, we profiled chromatin accessibility in 115 individuals from the Type 1 Diabetes Genetics Consortium (T1DGC), consisting of 57 controls and 58 T1D cases; 67 AFR and 48 EUR. We purified $CD4^+$ T cells from viably frozen PMBC samples via magnetic cell separation according to the manufacturer protocol using either negative selection (N=42; STEMCELL Technologies EasySep Human $CD4^+$ T Cell Isolation Kit) or positive selection (N=73; MACS Miltenyl Biotec). The selection approach was accounted for in subsequent data processing and analysis (see Online Methods "caQTL analysis"). After $CD4^+$ T cell purification, we followed the "Omni-ATAC-seq" protocol [26] for nuclei isolation, transposase incubation, and library preparation. Libraries were sequenced using 75 bp paired-end reads on an Illumina NextSeq.

**caQTL analysis**

Raw ATAC-seq data from patient samples were processed using the PEPATAC pipeline (http://code.databio.org/PEPATAC). Briefly, reads were trimmed using Skewer [33] and, after removing reads mapping to mitochondrial and human repeat regions, reads were mapped to GRCh38 using bowtie2 [34]. PCR duplicates were removed, enzymatic cut sites were inferred based on read alignment, and peaks were called using macs2 [29]. Libraries with transcription start site (TSS) enrichment scores less than 6 or fewer than 10 million aligned reads were excluded from analyses. A set of consensus peaks was determined by merging peaks across all samples using bedops [35]. Then, a matrix of peak counts was calculated by counting the number of cut sites within each consensus peak in each sample using the R package *bigWig* (https://github.com/andrelmartins/bigWig).

Peaks with low counts were excluded (required ≥ 10 reads in ≥ 50% of samples). Further peak quality filtering and normalization was performed using the *edgeR* [36] and *limma* R packages. These steps included:

(1) filtering for peaks with ≥ 10 counts-per-million (CPM) across samples within each batch;

(2) peak count normalization using the trimmed mean of M-values (TMM) method [37];

(3) mean-variance modeling-based transformation using the ‘voom’ function to enable linear modeling of peak counts assuming a normal distribution;

(4) removing outlier peaks by clustering samples based on counts for each peak (one at a time using k-means with k=2) and excluding any peak that results in one sample clustering separately from all other samples.

We confirmed matching sample identity between ATAC-seq libraries and genotyped subjects using the “Match BAM to VCF” (MBV) command in the software tool set QTLtools [38]. We tested for association between imputed genotype dosage and chromatin accessibility (caQTL analysis) by fitting a linear model, adjusting for the first two genotype principal components, age at sample collection, TSS enrichment score, and $CD4^+$ T cell purification approach using the R package *MatrixEQTL* [39]. The caQTL discovery analyses were done separately by ancestry group (EUR and AFR) and then combined in an inverse-variance weighted fixed effect meta-analysis with the R package *meta*. We tested all variant-peak combinations where the accessibility peak was within 1Mb of a T1D credible variant.

**Colocalization analysis**

We evaluated colocalization of T1D and caQTL for all peaks where at least one T1D credible variant (as defined by GUESSFM) was associated with peak accessibility with meta-analysis $p < 5\times10^{-5}$. We used the R package *coloc* [40] to formally test for colocalisation and the R package *locuscomparer* [41] to visualize colocalized signals. Because *coloc* assumes a single causal variant underlying both trait associations, we used conditional summary statistics in regions predicted to have more than one causal variant underlying the T1D association or regions with multiple, conditionally independent variants associated with accessibility of the same peak. When running *coloc* for T1D-caQTL colocalization, we used a prior probability of colocalisation of $5\times10^{-6}$, and provided association betas and standard errors as input data. When running *coloc* for T1D-eQTL colocalization, we used the same priors and supplied association z-scores. We considered GWAS and QTL signals to be significantly colocalized when the posterior probability of colocalisation was greater than 0.8 ('PP.H4.abf' > 0.8).

**Allele-specific accessibility analysis**

For significant caQTLs that colocalized with T1D-associated variants, we tested for allele-specific accessibility of the caQTL peak. First, we identified individuals heterozygous for T1D credible variants overlapping the caQTL peak itself. Within each heterozygous individual, we then counted the number of reads overlapping the variant position containing the reference or alternative allele. We only performed this analysis if the T1D credible variant overlapping the caQTL peak was directly genotyped on the ImmunoChip, since uncertainty in the heterozygous status of an individual could lead to biased results. For peaks with at least 5 participants who had at least 5 reads overlapping the peak, we formally tested whether the proportion of reads containing an alternative allele significantly deviated from the expected null hypothesis proportion of 0.5. We calculated p-values for deviation from "allelic balance" (proportion = 0.5 for each read) by fitting a generalized linear mixed model where the dependent variable is the number of reads and follows a Poisson distribution and the independent variables include a fixed effect for the allele and a random effect for the participant.

**Cell line for EMSA Supershift Assay**

Jurkat cell line (E6-1) purchased from ATCC and grown in Roswell Park Memorial Institute, RPMI (RPMI-1640; Gibco) supplemented with 10% fetal bovine serum, 1% penicillin-streptomycin, 1% sodium pyruvate), at 37 °C and 5% $CO_2$.

**EMSA Supershift Assay**

Labeled (5' IRDye 700) and unlabeled 31bp, single-stranded oligonucleotides containing rs72928038 were obtained from Integrated DNA Technologies (Reference Allele strand: 5'AGGGACGGATTTCCTGTAAGCTGATCTTGAA 3' and Alternative Allele strand: 5' AGGGACGGATTTCCTATAAGCTGATCTTGAA 3') along with complementary oligonucleotides. Double-stranded oligonucleotides were generated by annealing equal amount of labeled or unlabeled complementary oligonucleotides at 95 °C for 5 mins, followed by gradual cooling with a ramp rate of -1.2 °C/min for 1 hour (Bio-Rad C1000 Touch Thermal Cycler). Nuclear extract from Jurkat cells was obtained by following the manufacturer's protocol for NE-PER™ Nuclear and Cytoplasmic Extraction Reagents kit (Thermo Scientific) and the extracted nuclear protein was dialyzed with Slide-A-Lyzer MINI Dialysis Units, 10,000 MWCO (Thermo

Scientific) against a 1 L buffer (10 mM Tris, pH7.5, 50 mM KCl, 200 mM NaCl, 1 mM dithiothreitol, 1 mM phenylmethylsulfonyl fluoride, and 10% glycerol) for 16 h at 4 °C with slow stirring.

Binding reaction for the EMSA was carried out using 2 µL 10X binding buffer (100 mM Tris, 500 mM KCl, 10 mM DTT; pH 7.5), 2 µL 25 mM DTT (2.5% Tween 20), 1 µL Poly (dI-dC) (1 µg/µL in 10 mM Tris, 1 mM EDTA; pH 7.5), 1 µL 1% NP-40, 100 mM $MgCl_2$, 20 fmol IRDye double-stranded oligonucleotide probe, and 16 µg Jurkat nuclear extract in a final volume of 20 µL. For supershift lanes, tested transcription-factor-binding antibodies were diluted 1:50 with dd$H_2O$. Negative control Rabbit IgG was diluted to the same concentration as tested antibody. 1 µL of diluted antibody was added to the binding reaction mixture while maintaining a total volume of 20 ul. Binding reaction was incubated for 20 minutes at room temperature, after which 2 µL of 10X Orange Loading Dye was added. Electrophoresis was performed with binding reaction mixture on a pre-run 6% DNA retardation gel for 70 mins at 70V. To capture the image, the gel was placed directly on the Odyssey-CLx (Licor) scan bed. The gel was scanned with a thickness of 0.5 mm at 700 nm channel. The EMSA binding condition for rs72928038 was repeated three times to ensure reproducibility of the experiment.

**Priority Index (Pi)**

The data used to identify eQTL colocalization (eGenes) were the same as in the initial publication [42], namely unstimulated monocytes [43] (N =414), LPS stimulated monocytes after 2 hours [43] (N=261), LPS stimulated monocytes after 24 hours [43] (N =322), interferon-gamma stimulated monocytes after 24 hours [43] (N =367), unstimulated B cells [44] (N=286), unstimulated

NK cells (unpublished) (N =245), unstimulated neutrophils [45] (N =114), unstimulated $CD4^+$ T cells [46] (N =293), unstimulated CD8 T cells [46] (N =283), whole blood [47] (N =5,311). We also included eQTL data from a larger whole blood study (N =31,684) [24].

The data used to identify genes interacting with index variants (cGenes) were Hi-C data from monocytes, fetal thymus, naïve $CD4^+$ T cells, total $CD4^+$ T cells, activated total $CD4^+$ T cells, non-activated total $CD4^+$ T cells, naïve $CD8^+$ T cells, total $CD8^+$ T cells, naïve B cells, total B cells [48]. The data used to define functional genes (fGenes, pGenes and dGenes) were the same as those used in the original publication. Data from the STRING database [49] were used to define protein-protein interaction networks, where confidence scores ≥700 were considered.

**Data and code availability**

Code used to generate the results presented in this paper are available at https://github.com/ccrobertson/t1d-immunochip-2020. All univariable summary statistics for genotype association with T1D (including imputed variants) are available for download in **Supplementary Table 9**.
