## Supplementary Materials for "Fine-mapping, trans-ancestral and genomic analyses identify causal variants, cells, genes and drug targets for type 1 diabetes"

### Supplementary Figures

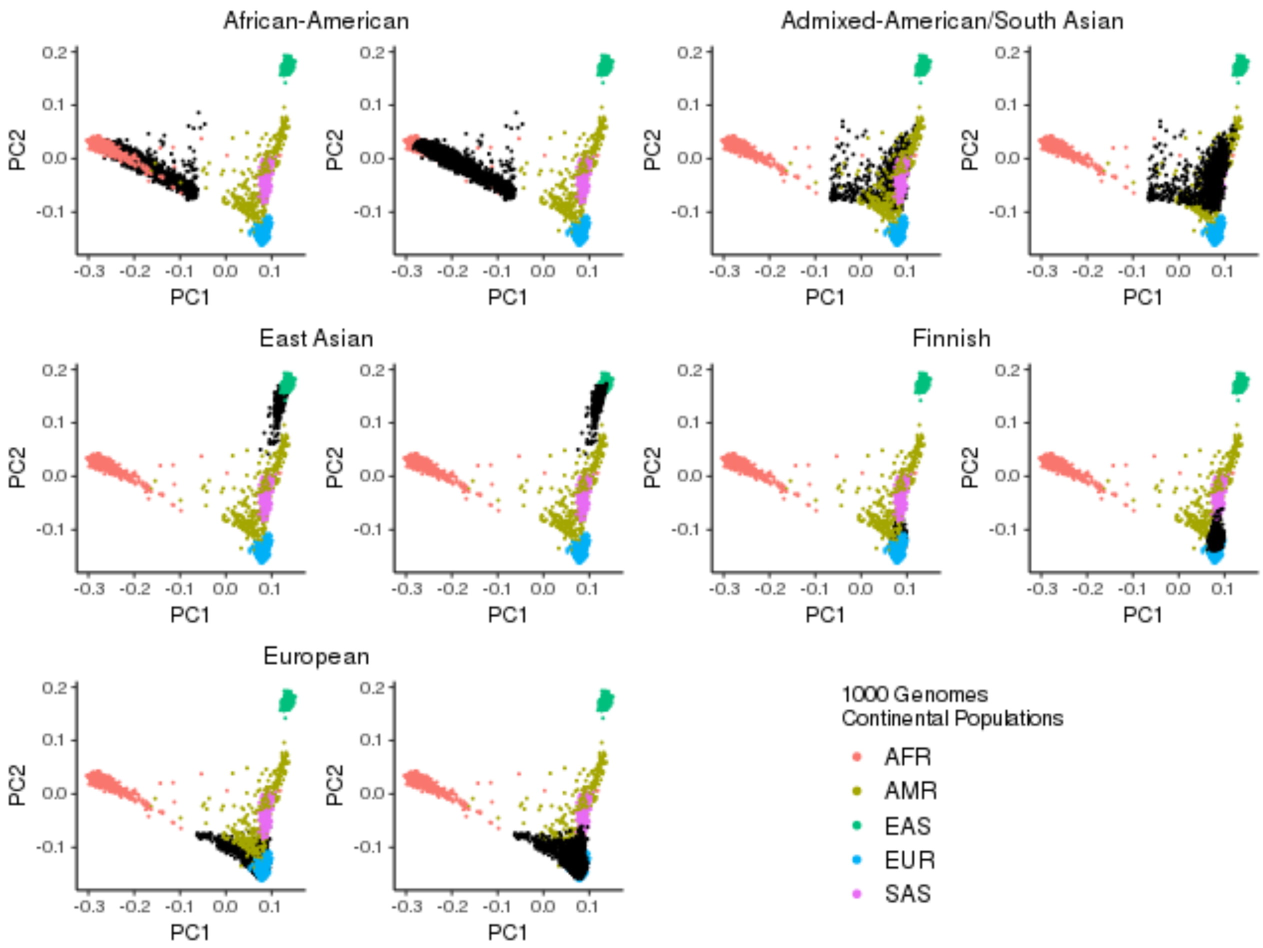

**Supplementary Figure 1:** Population-structure in the study population. The first versus second genetic principal components projected onto the 1000 Genomes Project principal components. Data in this study colored black, data from 1000 Genomes Project colored by ancestry group. Two plots shown for each ancestry group, one with the 1000 Genomes Project data at the front, the other with the data in this study at the front.

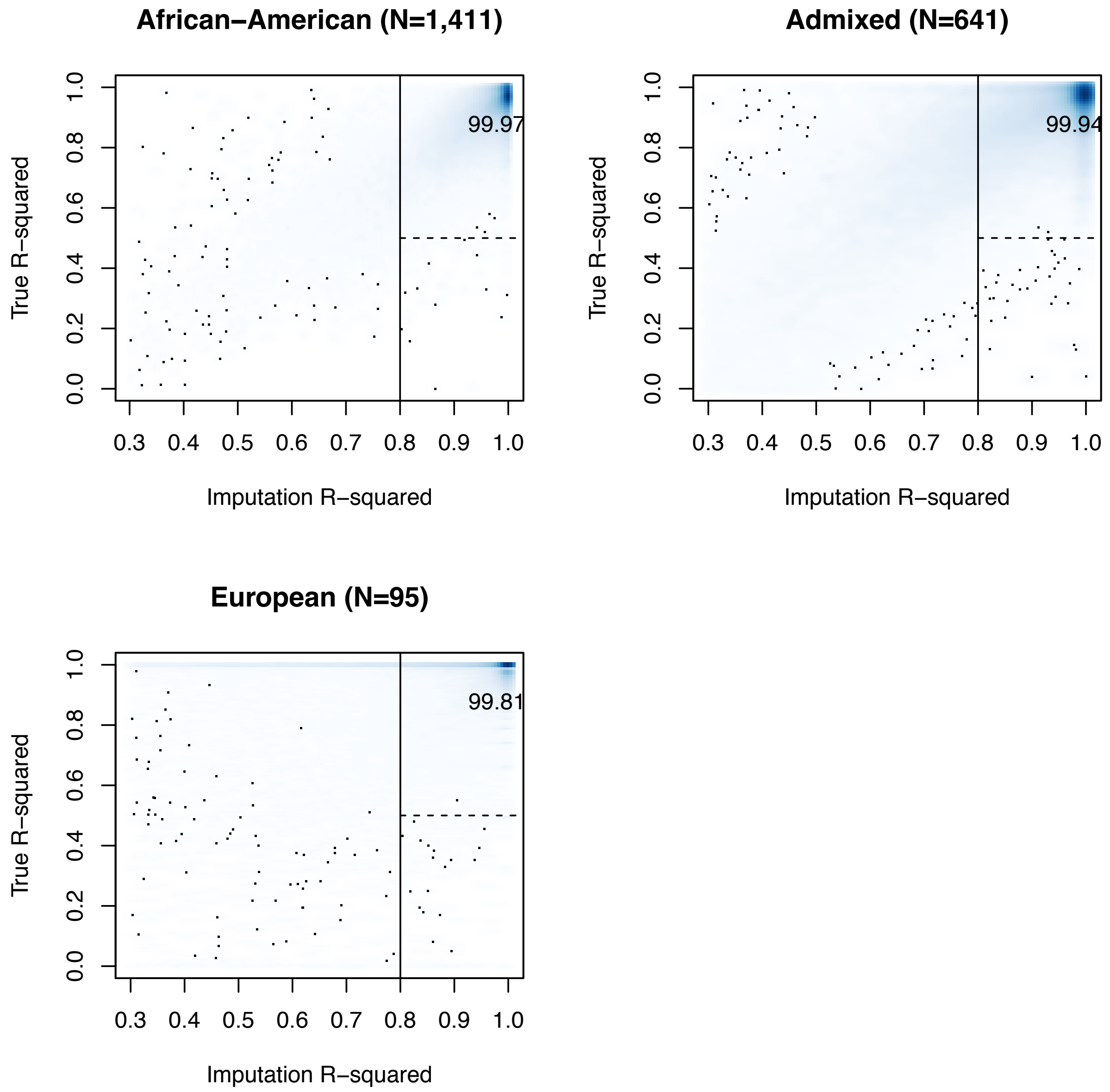

**Supplementary Figure 2:** Genotype accuracy for variants in ImmunoChip regions based on a subset of 2,147 participants with available whole genome sequence (WGS) data. "Imputation R-squared" is estimated imputation quality returned by the imputation software Minimac4. "True R-squared" is the Pearson correlation between genotypes obtained through imputation to the TOPMed reference panel versus WGS. Among variants with Imputation R-squared > 0.8 (right of solid vertical line), more than 90% have True R-squared > 0.5 in all three ancestry groups.

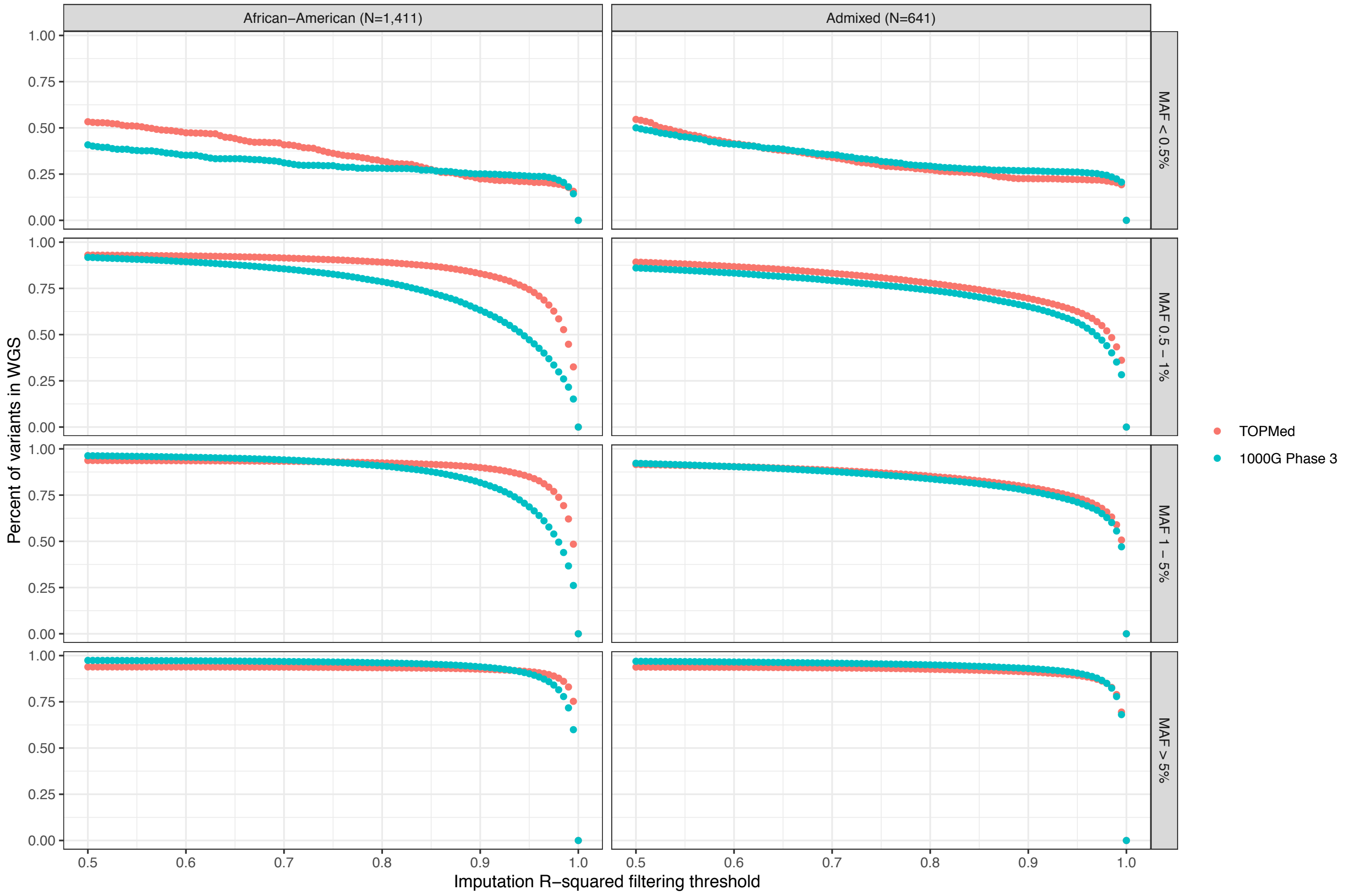

**Supplementary Figure 3:** Imputation coverage of ImmunoChip regions across a spectrum of imputation quality filtering thresholds and minor allele frequencies. Y-axis shows the proportion of variants detected by whole genome sequence (WGS) data that were imputed using the TOPMed (red) or 1000 Genomes Project Phase 3 (blue) reference panel. “Imputation R-squared” is estimated imputation quality returned by the imputation software Minimac4. MAF, minor allele frequency.

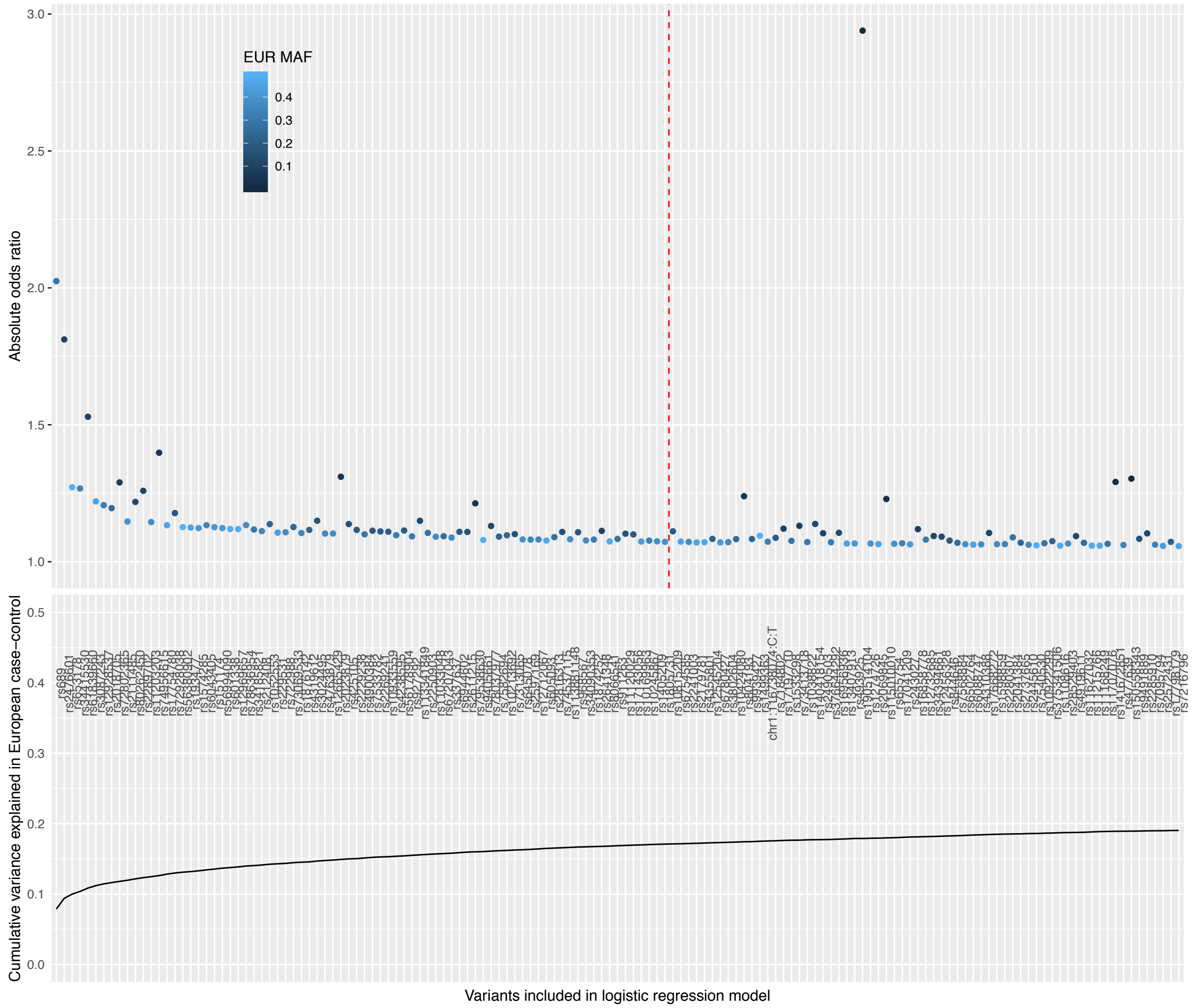

**Supplementary Figure 4:** Top panel: Absolute odds ratios for the lead variant in each T1D-associated region based on FDR < 0.01. Variants are coloured by minor allele frequency (MAF) in the European ancestry collection (lighter blue corresponding to higher MAF). Those to the left of the dashed line attained genome-wide significance ($p < 5\times10^{-8}$). Bottom Panel: Variance explained from logistic regression model using EUR case-control data only, from left to right, cumulatively adding variants to the logistic regression model; calculating the McFaddon's $r^2$ as a proxy for variance explained.

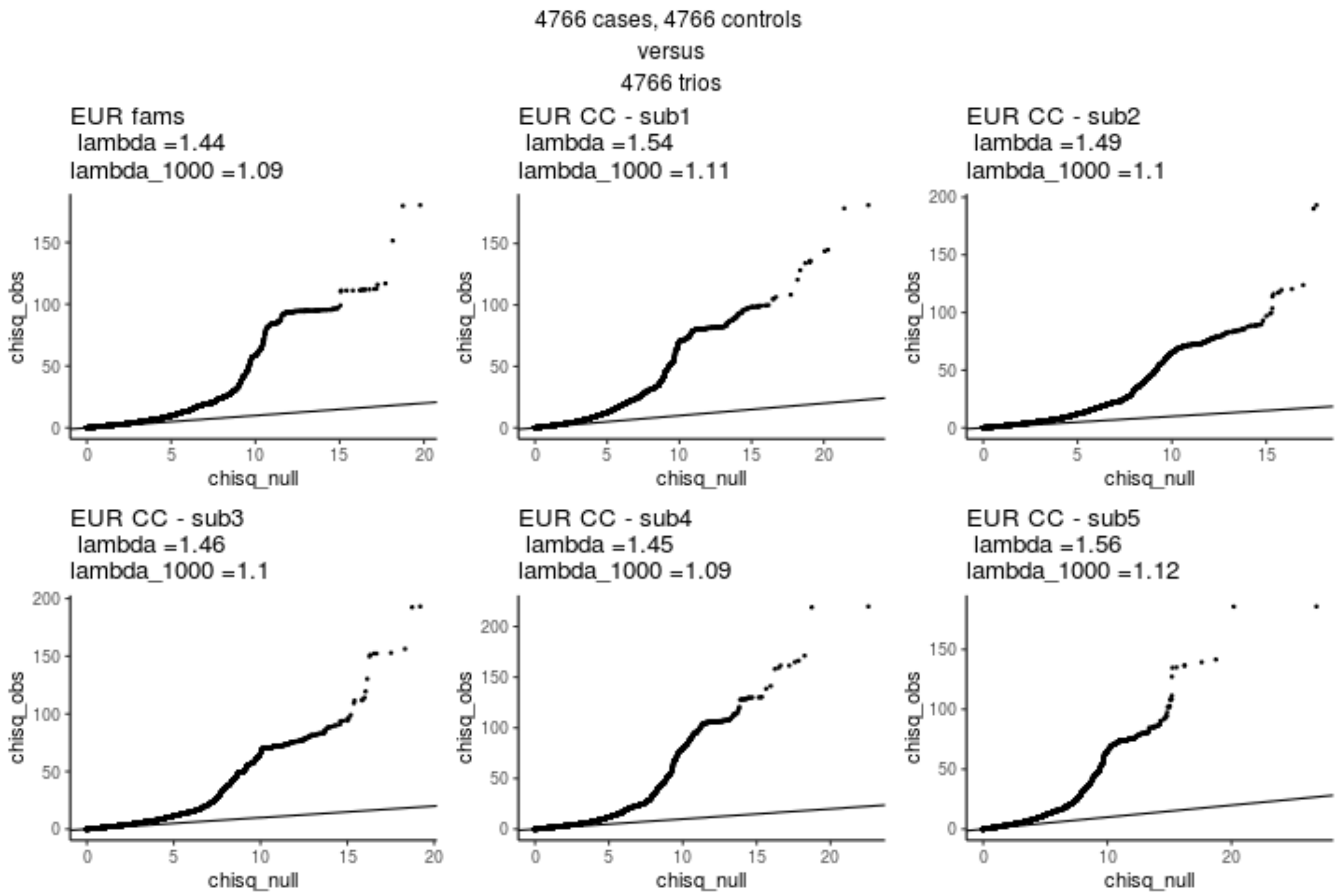

**Supplementary Figure 5:** Quantile-quantile plots showing the expected chi-square association statistics against the observed chi-square association statistics from the Phase II European family-based analysis results compared to five randomly sampled European case-control cohorts with equivalent statistical power to the family-based analysis.

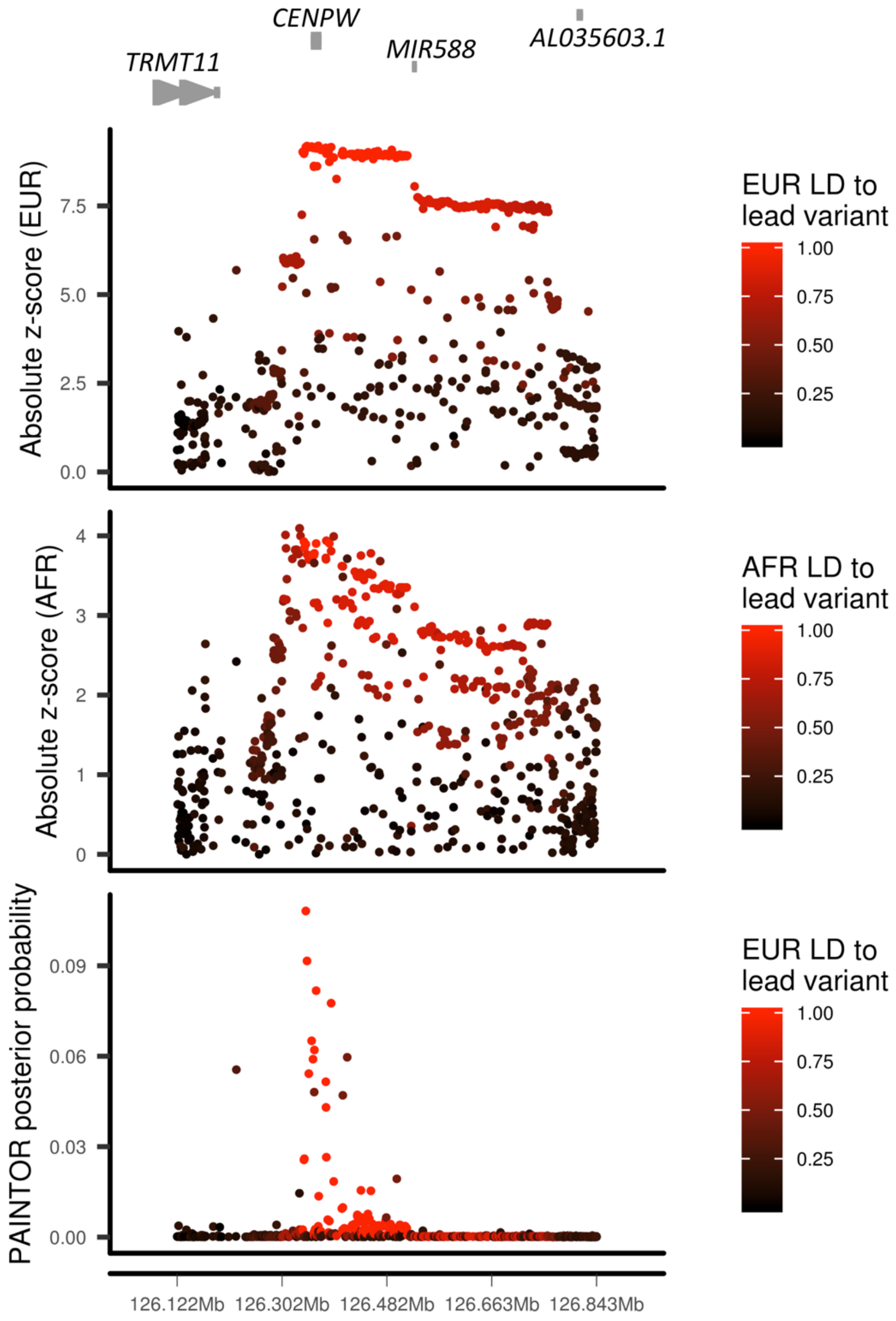

**Supplementary Figure 6:** Fine-mapping of the chromosome 6q22.32 region. European (EUR) and African (AFR) ancestry group association statistics, colored by ancestry group linkage disequilibrium (LD, lighter red corresponding to higher LD) to the lead PAINTOR-prioritised variant, of which the posterior probabilities are shown in the bottom panel.

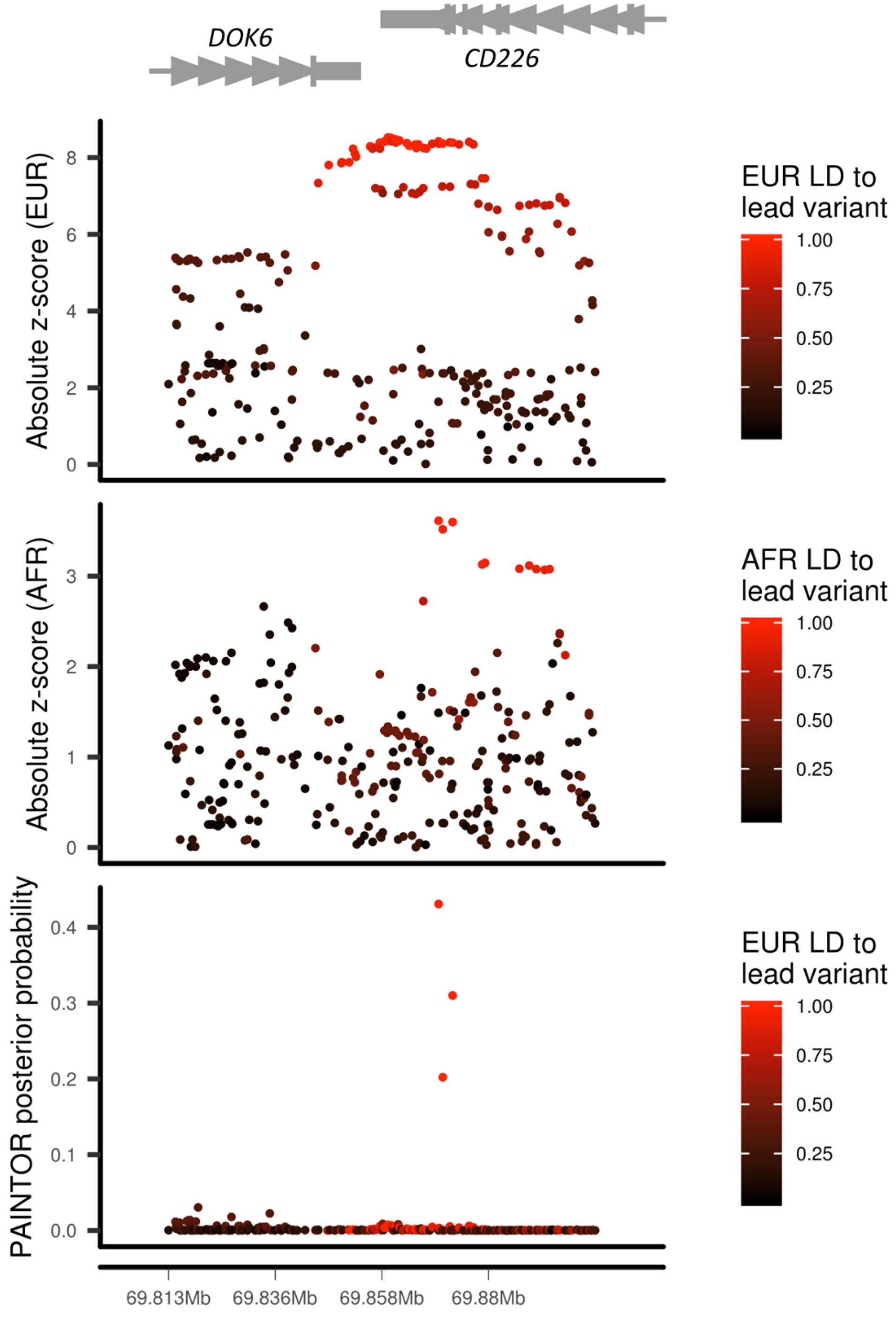

**Supplementary Figure 7:** Fine-mapping of the chromosome 18q22.2 region. European (EUR) and African (AFR) ancestry group association statistics, colored by ancestry group linkage disequilibrium (LD, lighter red corresponding to higher LD) to the lead PAINTOR-prioritised variant, of which the posterior probabilities are shown in the bottom panel.

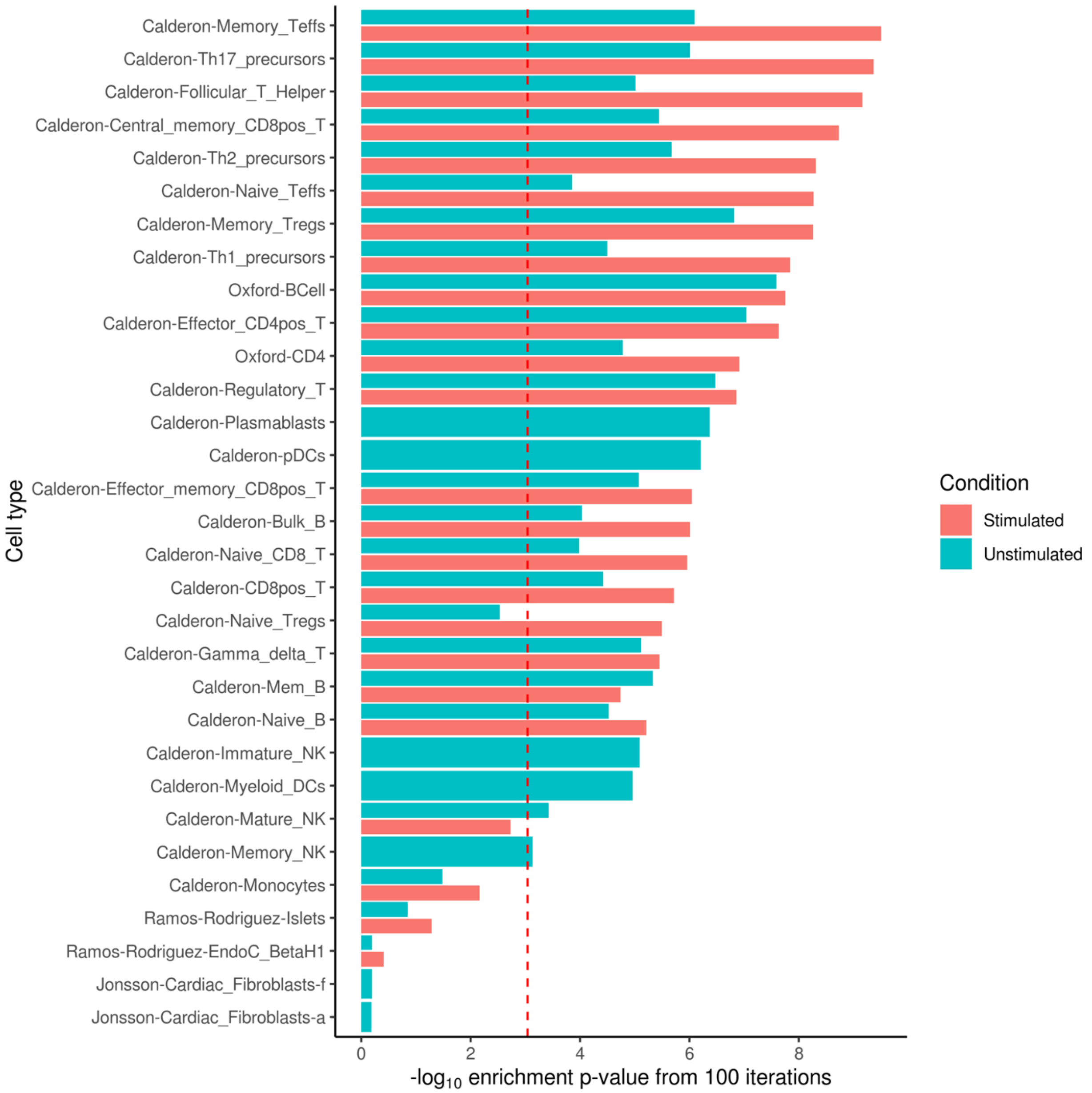

**Supplementary Figure 8:** Enrichment of T1D credible variants in ATAC-seq peaks in each cell type (red bars, stimulated; green bars, unstimulated), red dashed line represents the Bonferroni significance threshold at the 5% level.

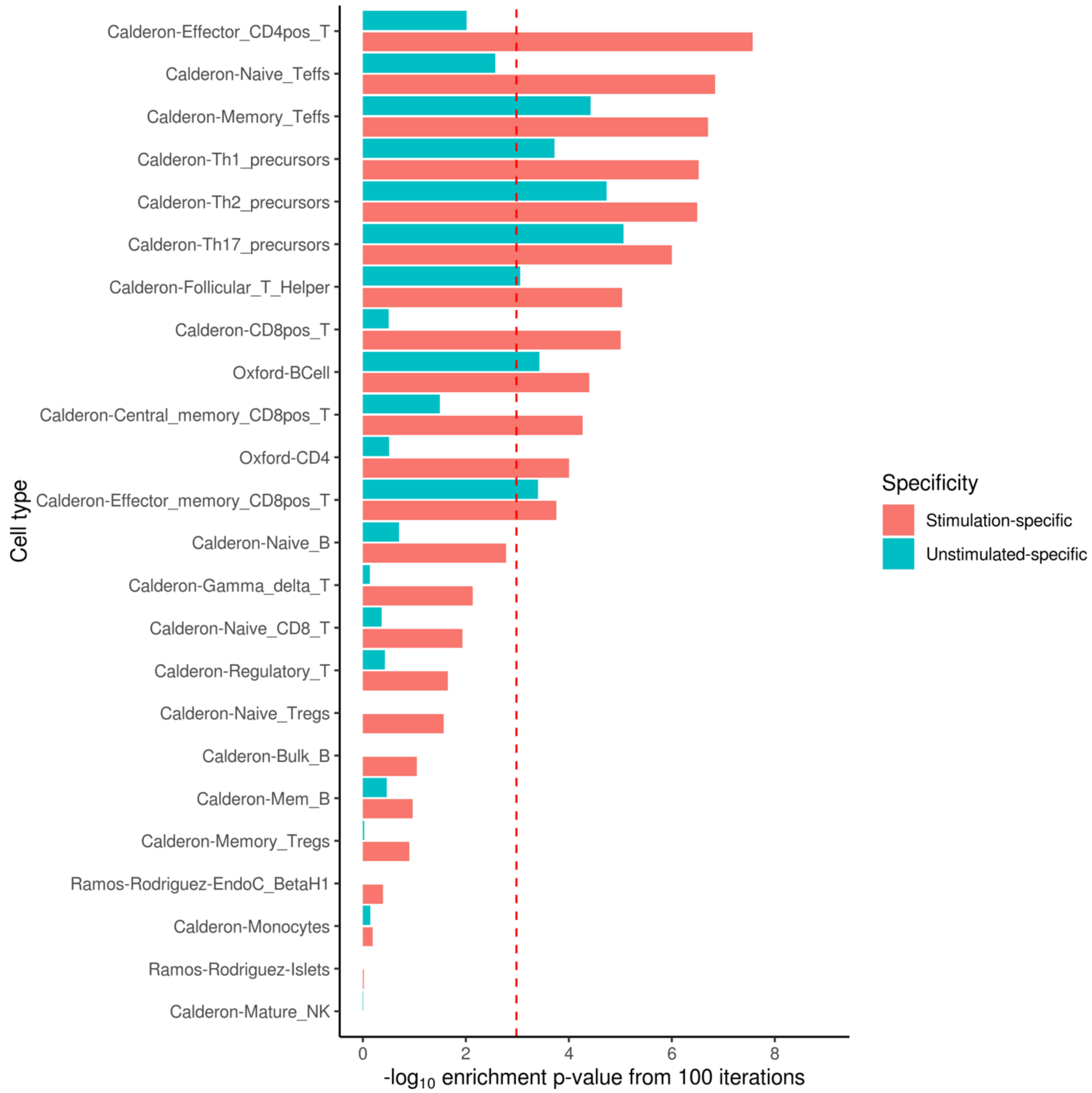

**Supplementary Figure 9:** Enrichment of T1D credible variants in differentially open ATAC-seq peaks between stimulation conditions, defined from a consensus list of peaks (Online Methods). Red bars show differentially open peaks in stimulated cells; green bars show differentially open peaks in unstimulated cells. Red dashed line is the Bonferroni significance threshold at the 5% level.

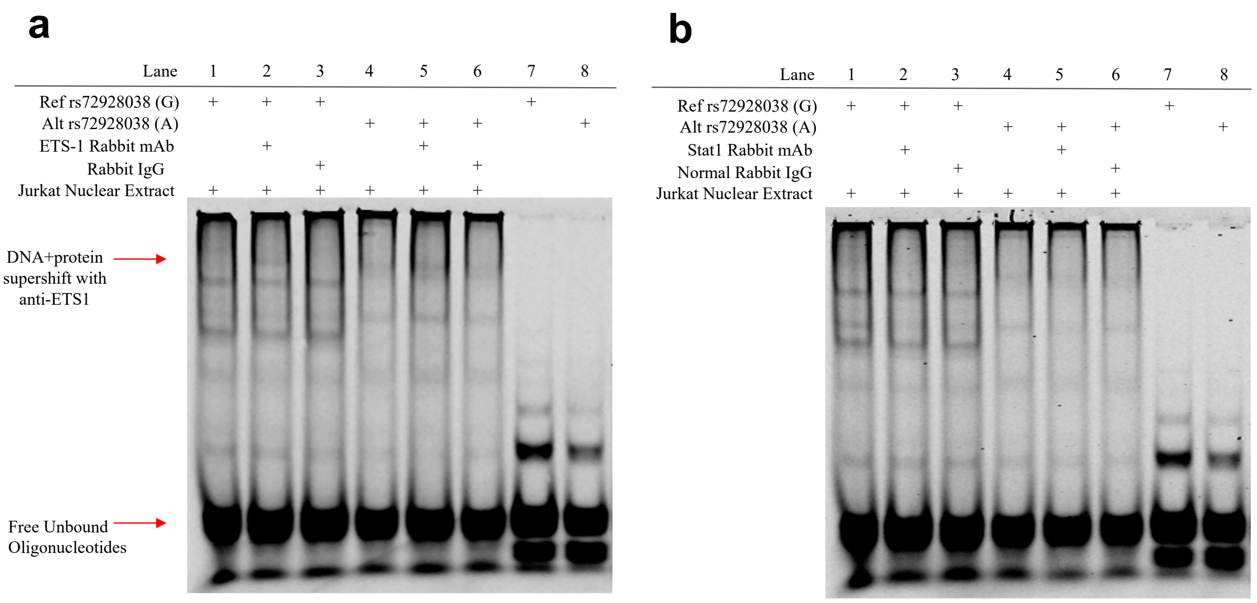

**Supplementary Figure 10: a)** rs72928038 with ETS-1 antibody supershift Electrophoretic Mobility Shift Assay (EMSA). The ETS-1 supershift EMSA demonstrates an allele specific supershift with rs72928038 G allele, while a shift is not observed with the A allele probe. Rabbit IgG was added to lane 3 and 6 as negative control for the supershift assay. **b)** rs72928038 with Stat1 Antibody Supershift EMSA Assay. Lane 1-3 and 7 contains the reference allele (G) of rs72928038 labeled probe. Lane 4-6 and 8 contains the alternative allele (A) of rs72928038 labeled probe. Lane 7 and 8 are negative controls. Rabbit IgG was added to lane 3 and 6 as negative controls for the supershift assay.

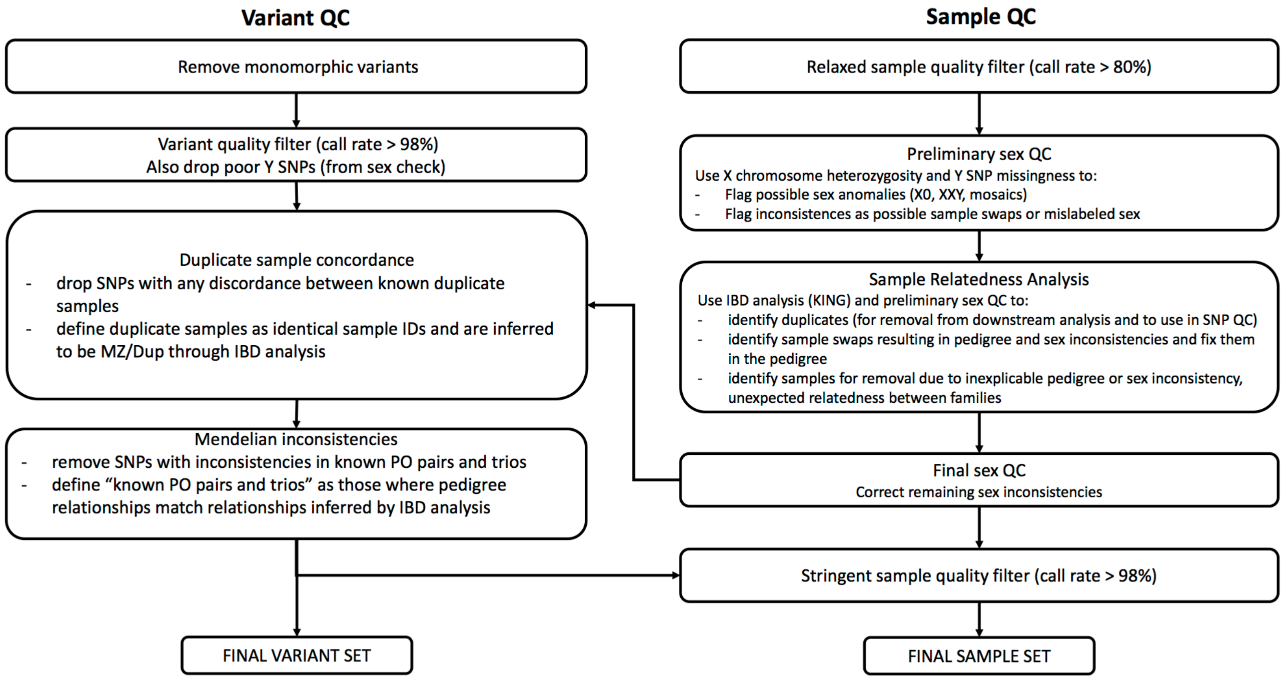

**Supplementary Figure 11:** Sample and variant quality control pipeline prior to imputation. See Online Methods and code repository (https://github.com/ccrobertson/t1d-immunochip-2020) for more details.

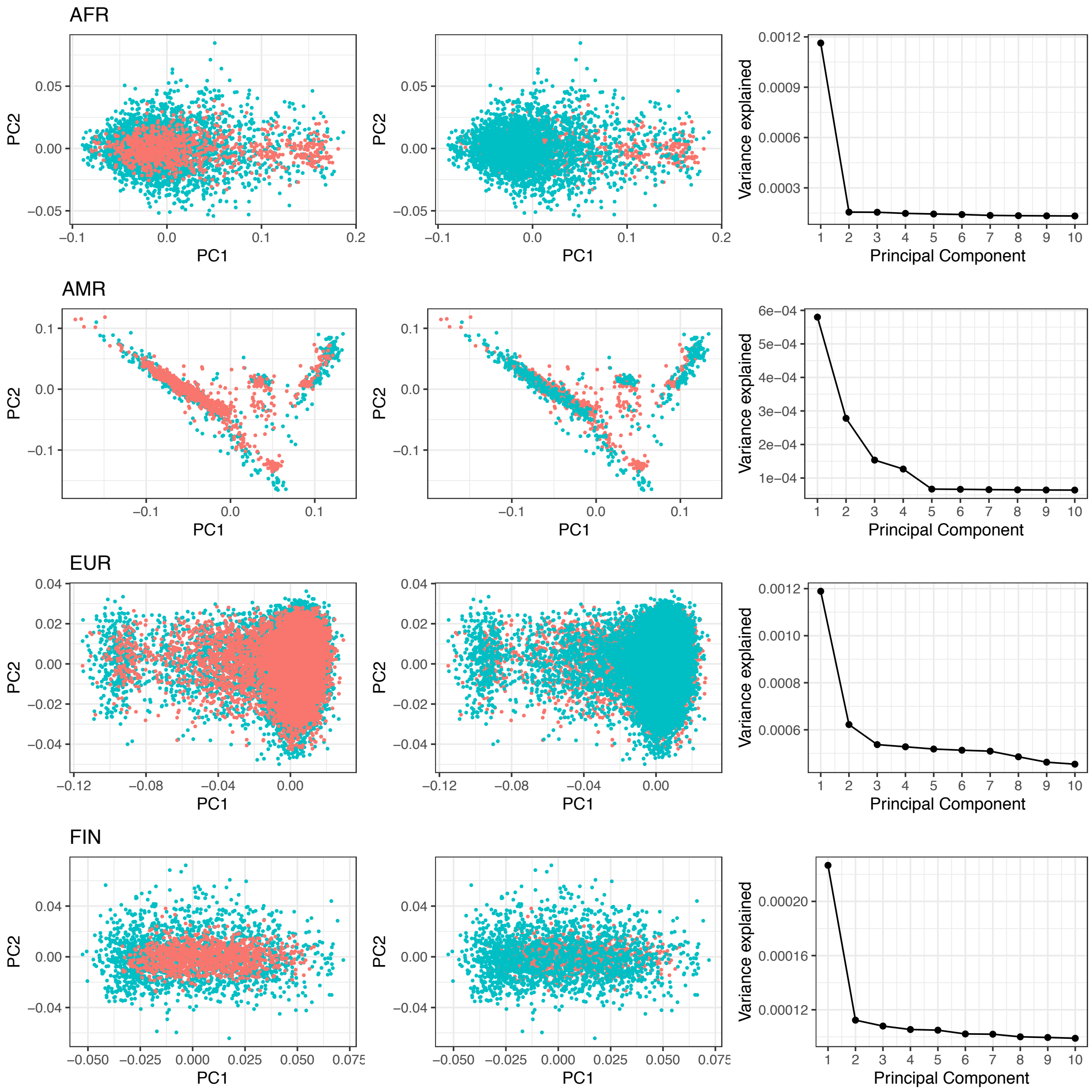

**Supplementary Figure 12:** Principal component analysis for cases (red) and controls (turquoise) for each ancestry group. Cases are plotted on top (left) or bottom (middle). Scree plots (right) suggest that linear models for genetic association include up to five principal components as covariates.

### Supplementary Note 1

Accuracy and coverage of imputation was assessed using WGS in 1,411 AFR, 641 AMR, and 95 EUR subjects. Specifically, as a measure of imputation accuracy, for each single nucleotide variants (SNV) we calculated the Pearson correlation coefficient and R-squared between genotypes obtained through imputation versus WGS. To measure imputation coverage of ImmunoChip regions, we calculated the proportion of SNVs with MAF >0.005 detected through WGS that were included in the imputed variant set after quality filtering at a range of imputation R-squared thresholds. Relative coverage of imputation based on TOPMed and 1000 Genomes reference panels was assessed in the AFR and AMR groups, where we had adequate number of samples with available WGS. After filtering for imputation R-squared >0.8, more than 99% of imputed SNVs within ImmunoChip regions were concordant with WGS with true R-squared >0.5 (**Supplementary Figure 2**). To quantify the coverage of ImmunoChip regions after imputation, we calculated the proportion of SNVs detected in WGS that were imputed with high confidence. Among 1,411 AFR and 641 AMR subjects, 92.3% and 87.6% of variants in ImmunoChip regions detected in WGS with MAF > 0.005 were imputed with imputation R-squared > 0.8, respectively (**Supplementary Figure 3**). Only variants within ImmunoChip regions or regions with relatively high variant density, defined as more than 50 variants genotyped in a 500kb region (**Supplementary Tables 2 and 3**), were included in the analysis, since the imputation of variants outside these regions would be based on a small number of genotyped variants only.

### Supplementary Note 2

Despite controlling for population stratification by analyzing major ancestry groups separately and adjusting for within-ancestry principal components in each ancestry-specific case-control analysis (Online Methods), the genomic inflation factors ($\lambda_{GC}$) from the complete meta-analysis was 1.40. Since the ImmunoChip intentionally covers regions of the genome previously associated with immune-mediated disease, $\lambda_{GC}$ for association with T1D across ImmunoChip variants is *a priori* anticipated to be greater than one. However, it is important to differentiate inflation due to enrichment of true biological association from inflation due to experimental artifact, such as population stratification. Due to the non-uniform distribution of ImmunoChip variants across the genome, LD-score regression (a common approach to determining sources of inflated test statistics in GWAS) cannot be applied to this data set. Thus, to rule out population stratification, we compared $\lambda_{GC}$ from TDT analysis, which is robust to population stratification, to $\lambda_{GC}$ from case-control analysis of comparable statistical power. Specifically, for each ancestry group, we generated five randomly sampled case-control data sets, each containing one case and one control for each trio, which results in equivalent statistical power (1). For example, in our European ancestry cohort, there were 4,766 trios. Thus, we subsampled, out of 13,458 European cases and 20,143 European controls, five data sets each containing 4,766 cases and 4,766 controls. After excluding the major histocompatibility complex (MHC), insulin (*INS*) and protein tyrosine phosphatase, non-receptor type 22 (*PTPN22*) regions, the $\lambda_{GC}$ for the European family-based analysis was 1.44, while the average $\lambda_{GC}$ from five randomly sampled case-control data sets of equivalent power was 1.50 (**Supplementary Figure 5**). Similar results are seen when only considering directly genotyped variants (**Supplementary Table 10**). Together, these data suggest that the inflation in the association analysis cannot be explained by population stratification in our study cohort. Thus, we believe the observed inflation in both case-control and family-based analyses is most likely due to enrichment for true association signal in ImmunoChip regions.

(1) Mcginnis, R., Shifman, S. & Darvasi, A. Power and Efficiency of the TDT and Case-Control Design for Association Scans. *Behav* **32**, 135–144 (2002).

### Appendix 1 – T1DGC Contributors

**Members of the Type 1 Diabetes Genetics Consortium:**

*Asia-Pacific Network*: Karen Alcantara, Tracey Baskerville, Nines Bautista, Eesh Bhatia, Francois Bonnici, Thomas Brodnicki, Pik To Cheung, Peter Colman, Andrew Cotterill, Jenny Couper, Kim Donaghue, Denise Li-Meng Goh, Len Harrison, Hiroshi Ikegami, Tim Jones, Khalid Abdul Kadir, Nor Azmi Kamaruddin, Uma Kanga, Alok Kanungo, Gurvinder Kaur, Tania Kelly, Yann-Jinn Lee, Margaret Lloyd, Kah Yin Loke, Amanda Loth, Narinder Mehra, Tony Merriman, Grant Morahan, Namid Munkhtuvshin, Araceli Panelo, Fraser Pirie, Niru Ratnam, C. B. Sanjeevi, Sandeep Sreedharan, Brian Tait, Allison Thomas, Jinny Willis, Sang Yanmei.

*European Network*: Francisco J. Ampudia-Blasco, Jesus Argente, Magdalena Avbelj, Gulja Babadjanova, Klaus Badenhoop, Lubomir Barak, Christos Bartsocas, Tadej Battelino, Emilia Belda, Polly Bingley, Bernhard O. Boehm, Ezio Bonifacio, Fatima Bosch, Eulalia Brugues, Raffaella Buzzetti, Joyce Carlson, Luis Castano, Anna Casu, Ondrej Cinek, Alberto de Leiva, Virginia Ruiz Esquide, Ana Fagulha, Marta Hernandez Garcia, Per-Henrik Groop, Cristian Guja, Alona Hamou, Erifili Hatziagelaki, Dan Headon, Simon Heath, Kaire Heilman, Robert Hermann, Nora Hosszufalusi, Lorenzo Iughetti, Cecile Julier, Ida Kinalska, Ingrid Kockum, Kalinka Koprivarova, Adam Kretowski, Dora Krikovszky, Nebojsa Lalic, Nicole Lambracht, Merce Lara, Mark Lathrop, Katharina Laubner, Ake Lernmark, Claire Levy-Marchal, Johnny Ludvigsson, Mara Marga, Antonio Martinez, Didac Mauricio, Nina Meier, Jorn Nerup, Antanas Norkus, Anna Okruszko, Carla Paganin, Xavier Palomer, Teresa Pedro, Moshe Phillip, Valdis Pirags, Flemming Pociot, Galina

Popova, Paolo Pozzilli, Jean-Francois Prud'Homme, Bart O. Roep, Ute Christine Rogner, Silke Rosinger, Ana Maria Varela Sande, Ilhan Satman, Edith Schober, Jochen Seufert, Jan Skrha, Gyula Soltesz, Giatgen Spinas, Juraj Stanik, Tanya Szendeffy, Erik Thorsby, Vallo Tillmann, Dag Undlien, Vaidotas Urbanavicius, Luciana Valente, Bart Van der Auwera, Andriani Vazeou-Gerasimidi, Dzilda Velickiene, Ana Wagner, Markus Walter, Alistair Williams, Lotte Albret Wissing, Miroslav Wurzburger, Anette-G Ziegler.

*North American Network*: Alan Aldrich, Marilyn Alford, Linda Amstutz, Mark Anderson, Beenu Aneja, Bonita Baker, Janice Bartos, Holly Baugh, Dorothy Becker, Christophe Benoist, Noureddine Berka, Kathleen Breen, Sonya Bridgeman, Patricia Cleary, Debbie Conboy, Patrick Concannon, Roberta Cook, Robert Couch, Lori Covell, Mark Daly, Jayne Danska, Larry Dolan, David Donaldson, Alessandro Doria, Janice Dorman, Angela Dove, Lee Ducat, George Eisenbarth, Henry Erlich, Pamela Fain, Rosanna Fiallo-Scharer, Lois Finney, Kenneth Gabbay, Gladys Gaillard-McBride, Terri Gammer, Daniel Geraghty, Soumitra Ghosh, Steven Gitelman, Nat Goodman, Gregory Goodwin, Jinko Graham, Carla Greenbaum, David Greenberg, William Hagopian, Mary Halvorson, John Hansen, Stephanie Higgins, Joel Hirschhorn, Kim Holmquist, Leroy Hood, Michelle Hull, Anhaita Jamula, Judith Johansen, Kevin Kaiserman, Fouad Kandeel, Francine Kaufman, Liz Langeland, Jean Lawrence, Nancy Lewis, Victoria Magnuson, Jennifer Marks, Andrea Martin, Della Matheson, Beth Mayer-Davis, Marli McCulloh-Olson, Richard McIndoe, Brad McNeney, Eric Mickelson, Priscilla Moonsamy, Antoinette Moran, Patricia Mueller, Mary Murray, Gerald Nepom, David Ng, Janelle Noble, Jill Norris, Tihamer Orban, David Owerbach, Andrew Paterson, Helen Patrie, Diana B. Petitti, Catherine Pihoker, Constantin Polychronakos, Donna Prokopczak, Alberto Pugliese, Beeca Pyle, Philip Raskin, Natasha

Razack, Marian Rewers, Karen Riley, Henry Rodriquez, John Rogus, Ami Romanowski, Jerry Rotter, Monique Roy, Penny Satterwhite, Desmond Schatz, Gary Schoch, Mara Semel, Jin-Xiong She, Terry Smith, Janice Sowinski, Richard Spielman, Debbie Standiford, Caroline Suh, Christine Tam, Kent Taylor, Chrystal Thomas, Joan Thomas, Jay Tischfield, Ellen Toth, Deborah Truell, Diane Wherrett, Michelle Whiting, Theodora Wilson, Darrell Wilson, Lue Ping Zhao.

*United Kingdom Network*: Francesco Cucca, David Dunger, Graham Alec Hitman, Simon Howell, Sarah Nutland, Helen Rance, Luc Smink, John Todd, Jaakko Tuomilehto, Neil Walker, Barry Widmer, Heather Withers.

*Latin America*: Arturo Alvarado, Cresio Alves, Pablo Aschner, Elena Carrasco, Martha de Sereday, Laerico Franco, Teresa Frazer, Gustavo Frechtel, Clara Gorodezky, Jorge Jiminez, Ana Maria Jorge, Roberto Lanes, Ingrid Libman, Leonardo Mancillas, Carmen Mazza, Miguel Pasquel, Francisco Perez, Francisco Gomez Perez, Carmen Pisciottano, Ivelise Ramos, Olga Ramos, Maria Isabel Rojas, Antonio Selman-Geara, Gladys Veray.

*Coordinating Center*: Don Babcock, Stephanie Beck, Mark Brown, Cralen Davis, Mark Espeland, Mark Hall, Teresa Harnish, Laura Hemrick, Joan Hilner, Letitia Howard, Ethan M. Lange, Carl Langefeld, Josyf Mychaleckyj, June Pierce, David Reboussin, Stephen Rich, Scott Rushing, Michele Sale, Elizabeth Sides, Michael Steffes, Augy Thiel, Lynne Wagenknecht, Dustin Williams, Jianzhao Xu.

### Appendix 2 – SEARCH Contributors

**SEARCH AUTHORSHIP LIST**
**(Listed within each site/center: PI(s) & then alphabetically within each institution/facility.)**

The writing group for this manuscript wishes to acknowledge the contributions of the following individuals to the SEARCH for Diabetes in Youth Study:

**SEARCH SITES**

***California:*** **(PI)** Jean M. Lawrence, ScD, MPH, MSSA
Peggy Hung, MPH; Corinna Koebnick, PhD, MSc; Xia Li, MS; Eva Lustigova, MPH; Kristi Reynolds, PhD, MPH for the Department of Research & Evaluation, Kaiser Permanente Southern California, Pasadena California, and David J. Pettitt, MD, Santa Barbara, California.

***Carolinas:*** **(PI)** Elizabeth J. Mayer-Davis, PhD
Amy Mottl, MD, MPH; Joan Thomas MS, RD for the University of North Carolina, Chapel Hill.

Malaka Jackson, MD; Lisa Knight, MD; Angela D. Liese, PhD, MPH; Christine Turley, MD for the University of South Carolina.

Deborah Bowlby, MD for the Medical University of South Carolina.

James Amrhein, MD; Elaine Apperson, MD; Bryce Nelson, MD for Greenville Health System and Eau Claire Cooperative Health Center.

***Colorado:*** **(PI)** Dana Dabelea, MD, PhD
Anna Bellatorre, PhD; Tessa Crume, PhD, MSPH; Richard F. Hamman, MD, DrPH; Katherine A. Sauder, PhD; Allison Shapiro, PhD, MPH; Lisa Testaverde, MS for the LEAD Center in the Department of Epidemiology, Colorado School of Public Health, University of Colorado Denver.

Georgeanna J. Klingensmith, MD; David Maahs, MD; Marian J. Rewers, MD, PhD; Paul Wadwa, MD for the Barbara Davis Center for Childhood Diabetes.

Stephen Daniels, MD, PhD; Michael G. Kahn, MD, PhD; Greta Wilkening, PsyD for the Department of Pediatrics and Children's Hospital.

Clifford A. Bloch, MD for the Pediatric Endocrine Associates.

Jeffrey Powell, MD, MPH for the Shiprock Service Unit, Navajo Area Indian Health Service.

Kathy Love-Osborne, MD for the Denver Health and Hospital Authority.

Diana C. Hu, MD for the Pediatrics Department, Tuba City Regional Health Care Center, Tuba City, AZ.

***Ohio*: (PI)** Lawrence M. Dolan, MD
Amy S. Shah, MD, MS; Debra A. Standiford, MSN, CNP; Elaine M. Urbina, MD, MS for the Cincinnati Children's Hospital Medical Center, Department of Pediatrics, University of Cincinnati.

***Washington*: (PI)** Catherine Pihoker, MD
Irl Hirsch, MD; Grace Kim, MD; Faisal Malik, MD, MSHS; Lina Merjaneh, MD; Alissa Roberts, MD; Craig Taplin, MD; Joyce Yi-Frazier, PhD for the University of Washington.

Natalie Beauregard, BA; Cordelia Franklin, BS; Carlo Gangan, BA; Sue Kearns, RN; Mary Klingsheim, RN; Beth Loots, MPH, MSW; Michael Pascual, BA for Seattle Children's Hospital.

Carla Greenbaum, MD for Benaroya Research Institute.

**CENTERS and LAB**

***Centers for Disease Control and Prevention*:** Giuseppina Imperatore, MD, PhD, Sharon H. Saydah, PhD

***National Institute of Diabetes and Digestive and Kidney Diseases, NIH*:** Barbara Linder, MD, PhD

***Central Laboratory:* (PI)** Santica M. Marcovina, PhD, ScD (PI)
Alan Chait, MD; Noemie Clouet-Foraison, PhD; Jessica Harting; Greg Strylewicz, PhD for the University of Washington Northwest Lipid Metabolism and Diabetes Research Research Laboratories.

***Coordinating Center*: (Co-PIs)** Ralph D'Agostino, Jr., PhD, Elizabeth T. Jensen, MPH, PhD; Lynne E. Wagenknecht, DrPH;
Ramon Casanova, PhD; Jasmin Divers, PhD; Maureen T. Goldstein, BA; Leora Henkin, MPH, M.Ed; Scott Isom, MS; Kristin Lenoir, MPH; June Pierce, AB; Beth Reboussin, PhD; Joseph Rigdon, PhD; Andrew Michael South, MD, MS; Jeanette Stafford, MS; Cynthia Suerken, MS; Brian Wells, MD, PhD; Carrie Williams, MA, CCRP for Wake Forest School of Medicine.
